# Quantitative firing pattern phenotyping of hippocampal neuron types

**DOI:** 10.1101/212084

**Authors:** Alexander O. Komendantov, Siva Venkadesh, Christopher L. Rees, Diek W. Wheeler, David J. Hamilton, Giorgio A. Ascoli

## Abstract

Systematically organizing the anatomical, molecular, and physiological properties of cortical neurons is important for understanding their computational functions. Hippocampome.org defines 122 neuron types in the rodent hippocampal formation (dentate gyrus, CA3, CA2, CA1, subiculum, and entorhinal cortex) based on their somatic, axonal, and dendritic locations, putative excitatory/inhibitory outputs, molecular marker expression, and biophysical properties such as time constant and input resistance. Here we augment the electrophysiological data of this knowledge base by collecting, quantifying, and analyzing the firing responses to depolarizing current injections for every hippocampal neuron type from available published experiments. We designed and implemented objective protocols to classify firing patterns based on both transient and steady-state activity. Specifically, we identified 5 transients (delay, adapting spiking, rapidly adapting spiking, transient stuttering, and transient slow-wave bursting) and 4 steady states (non-adapting spiking, persistent stuttering, persistent slow-wave bursting, and silence). By characterizing the set of all firing responses reported for hippocampal neurons, this automated classification approach revealed 9 unique families of firing pattern phenotypes while distinguishing potential new neuronal subtypes. Several novel statistical associations also emerged between firing responses and other electrophysiological properties, morphological features, and molecular marker expression. The firing pattern parameters, complete experimental conditions (including solution and stimulus details), digitized spike times, exact reference to the original empirical evidence, and analysis scripts are released open-source through Hippocampome.org for all neuron types, greatly enhancing the existing search and browse capabilities. This information, collated online in human-and machine-accessible form, will help design and interpret both experiments and hippocampal model simulations.

**Significance Statement:** Comprehensive characterization of nerve cells is essential for understanding signal processing in biological neuronal networks. Firing patterns are important identification characteristics of neurons and play crucial roles in information coding in neural systems. Building upon the comprehensive knowledge base Hippocampome.org, we developed and implemented automated protocols to classify all known firing responses exhibited by each neuron type of the rodent hippocampus based on analysis of transient and steady-state activity. This approach identified the most distinguishing elements of every firing phenotype and revealed previously unnoticed statistical associations of firing responses with other electrophysiological, morphological, and molecular properties. The resulting data, freely released online, constitute a powerful resource for designing and interpreting experiments as well as developing and testing hippocampal models.

## Introduction

Quantitative characterization of neurons is essential for understanding the functions of neuronal networks at different hierarchical levels. The hippocampus provides an excellent test-bed for this exploration as it is one of the most intensively studied parts of the mammalian brain, and is involved in critical functions including learning (Rudy and Sutherland 1995, 1989), memory (Eichenbaum et al., 1992; Eichenbaum, 2000, 2017), spatial navigation (Hafting et al. 2005; O’Keefe and Dostrovsky, 1971), and emotional associations (Buchanan, 2007).

Transmission of information between neurons is carried out by sequences of spikes, and firing rates are commonly believed to represent the intensity of input stimuli. Since the first discovery in sensory neurons (Adrian and Zotterman, 1926), this principle was generalized and extended to neurons from different brain regions including the hippocampus (McNaughton et al, 1983). However, it was also found that the firing rate of certain neurons is not constant over time even if the stimulus is permanently applied. One form of such time-dependent responses is spike frequency adaptation, manifested in a decrease of firing rate (Adrian and Zotterman, 1926). Neurons can produce diverse firing patterns in response to similar stimuli due to the inhomogeneity in their intrinsic properties (Connors and Gutnick, 1990). Both firing rates and temporal firing patterns are now recognized to play important roles in neural information coding (Ferster and Spruston, 1995).

In electrophysiological experiments *in vitro*, hippocampal neurons demonstrate a vast diversity of firing patterns in response to depolarizing current injections. These patterns are referred to by many names, including delayed, adapting, accommodating, interrupted spiking, stuttering, and bursting (Canto and Witter 2012a,b; Hemond et al., 2008; Lübke et al, 1998; Mercer et al., 2007; Pawelzik et al., 2002; Tricoire et al., 2011). Uncertainties and ambiguities in classification and naming of neuronal firing patterns are similar to other widely spread terminological inconsistencies in the neuroscience literature, posing obstacles to effective communication within and across fields (Hamilton et al., 2017).

Recent efforts aimed to develop firing pattern classification for identifying distinct electrical types of cortical neurons (Markram et al., 2004, 2015; Petilla Interneuron Nomenclature Group et al., 2008). Notably, statistical analysis of a large set of electrical features of neocortical interneurons with different firing patterns from a single lab yielded a refinement of the physiological component of the Petilla Nomenclature (Druckmann et al., 2013). However, comparisons across labs and experimental studies are typically limited to qualitative assessments of the illustrated firing traces or subjectively intuitive criteria. Moreover, firing pattern data are seldom unambiguously linked to neuron types independently defined by morphological and molecular criteria.

The Hippocampome.org knowledge base defines neuron types based on the locations of their axons, dendrites, and somata across 26 parcels of the rodent hippocampal formation, putative excitatory/inhibitory output, synaptic selectivity, and major and aligned differences in molecular marker expressions and biophysical properties (Wheeler et al., 2015). Version 1.3 of Hippocampome.org identified 122 neuron types in 6 major areas: 18 in dentate gyrus (DG), 25 in CA3, 5 in CA2, 40 in CA1, 3 in subiculum (SUB), and 31 in entorhinal cortex (EC). The core assumption of this identification scheme is that neurons with qualitatively different axonal or dendritic patterns, or with multiple substantial differences in other dimensions, belong to different types. For the majority of neuron types, Hippocampome.org reports 10 basic biophysical parameters that numerically characterize passive and spike properties (hippocampome.org/ephys-defs), consistent with other literature-based neuroinformatics efforts (Tripathy et al., 2015).

Here, we developed an objective numerical protocol to automatically classify all published electrophysiological recordings of somatic spiking activity for morphologically identified hippocampal neurons from Hippocampome.org. This process revealed specific firing pattern phenotypes, potential neuronal subtypes, and statistical associations between firing responses and other properties. Inclusion of the classified firing patterns and their quantitative parameters, along with a comprehensive tabulation of the underlying experimental conditions, substantially extends the online search and browse functionalities of Hippocampome.org, providing a wealth of annotated data for quantitative analysis and modeling.

## Materials and Methods

### Data collection, extraction and digitization

The firing patterns of hippocampal neurons were classified based on their spiking responses to supra-threshold step-current pulses of different amplitude and duration as reported in peer-reviewed publications. Firing pattern parameters were extracted from electronic figures using Plot Digitizer (plotdigitizer.sourceforge.net) for all Hippocampome.org neuron types (Wheeler et al., 2015) for which they were available (90 out of 122). A total of 247 traces were analyzed. We extracted values of first spike latency (i.e. delay), inter-spike intervals (ISIs), and post-firing silence (in ms), as well as slow-wave amplitude (in mV) for burst firing recording. For firing pattern identification and analysis, ISIs in each recording were normalized to the shortest inter-spike interval (ISI_min_) within that time series, to allow meaningful comparison.

All analyzed recordings were obtained in normal artificial cerebrospinal fluids (ACSF) from rodents (rats 85%, mice 12%, and guinea pigs 3%) generally described as ‘young adults’ (ages ranging from 11 to 70 days for rats and from 10 to 56 days for mice). All firing traces considered in this report were recorded in slice preparations; 74% of electrophysiological traces were obtained using whole-cell patch clamp and 26% intracellular recording with sharp microelectrodes. All experimental conditions and solution compositions were extracted and stored with every recording and are available at Hippocampome.org as specified in the “Web portal” section below. Representative examples of ACSF and of solutions for pipette filling are shown in Table 1 and Table 2, respectively.

**Table 1.** Representative examples of artificial cerebrospinal fluids

Illustrations and Tables
| NaCl (mM) | KCl (mM) | CaCl<sub>2</sub> (mM) | NaH<sub>2</sub>PO<sub>4</sub> (mM) | KH<sub>2</sub>PO<sub>4</sub> (mM) | MgCl<sub>2</sub> (mM) | MgSO<sub>4</sub> (mM) | NaHCO<sub>3</sub> (mM) | Glucose (mM) | t (° C) | Pct. | Sample<br>Reference |
| --- | --- | --- | --- | --- | --- | --- | --- | --- | --- | --- | --- |
| 126 | 3 | 2 | 1.25 |  |  | 2 | 26 | 10 | 34 | 30.0<br>% | Canto and Witter,<br>2012a |
| 124 | 3.3 | 2.5 |  | 1.2 |  | 1 | 25.5 | 15 | 34-35 | 11.3<br>% | Ali and Thomson,<br>1998 |
| 120 | 3.3 | 1.8 | 1.23 |  |  | 1.2 | 25 | 10 | 32-34 | 6.9% | Mott et al, 1997 |
| 130 | 3.5 | 2.5 | 1.25 |  | 1.5 |  | 24 | 10 | 32-34 | 6.5% | Tricoire et al, 2011 |
| 126 | 2.5 | 2 | 1.25 |  | 2 |  | 26 | 10 | 34-37 | 6.5% | Zemankovics et<br>al., 2010 |
| 126 | 3 | 2 | 1.25 |  |  | 2 | 24 | 10 | 34-35 | 5.3% | Buhl et al., 1994 |
| 125 | 2.5 | 2 | 1.25 |  | 1 |  | 25 | 25 | 35-37 | 4.5% | Lübke et al., 1998 |
| 124 | 5 | 2 | 1.25 |  |  | 2 | 26 | 10 | 33-35 | 2.8% | Hamam et al.,<br>2002 |
| 124 | 3 | 2.5 | 1.23 |  |  | 1.2 | 26 | 10 | 30 | 2.0% | Williams et al.,<br>2007 |

**Table 2.**
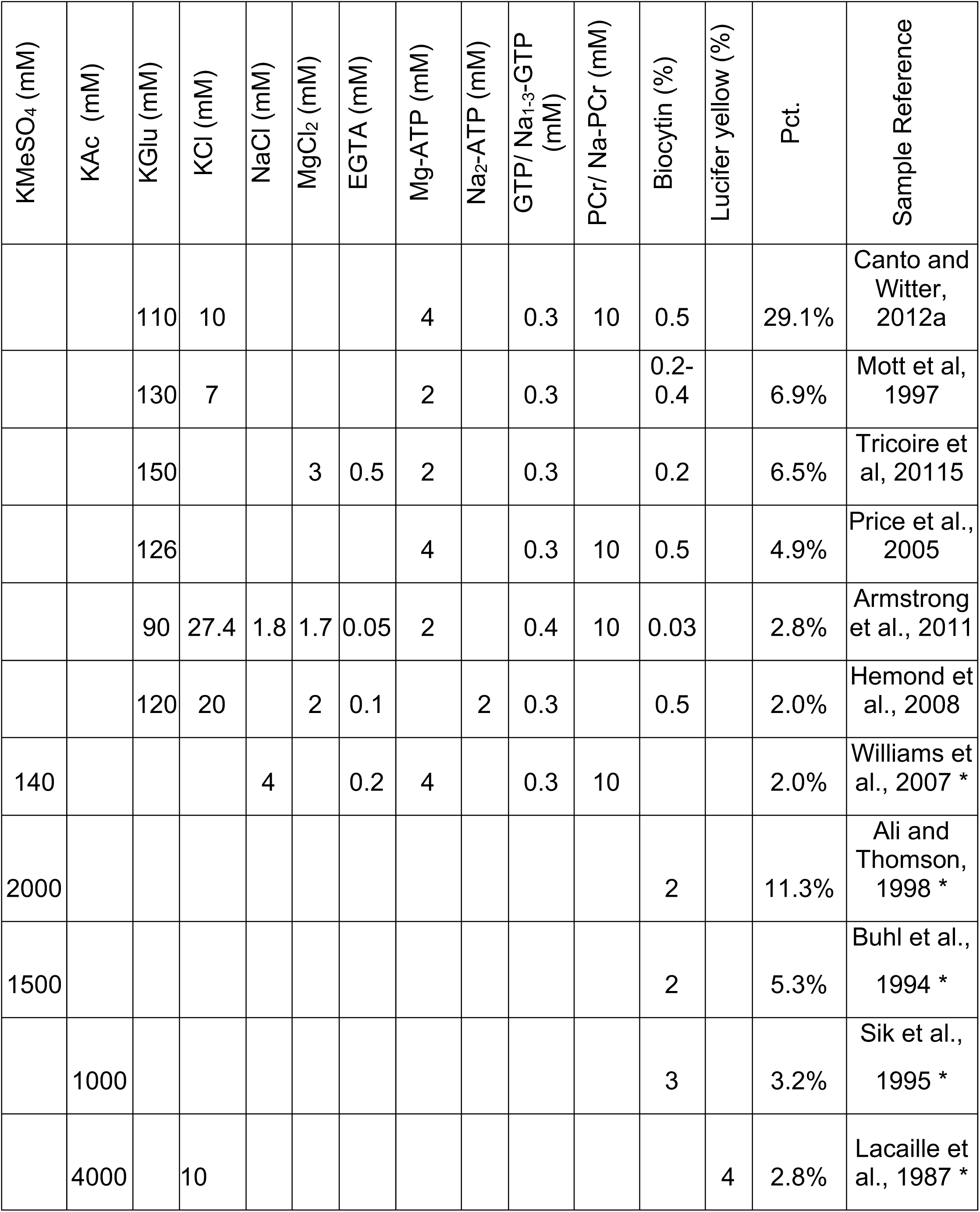
Representative examples of solutions for patch pipette and micropipette filling

### Firing pattern classification

Hippocampal neuron types display a variety of firing pattern elements in both their transient and steady state responses to continuous stimulation (Figure 1). Specifically, transients (which we label by dot-notation) can be visually differentiated into delay (D.), adapting spiking (ASP.), rapidly adapting spiking (RASP.), transient stuttering (TSTUT.), and transient slow-wave bursting (TSWB.). Steady states include silence (SLN), non-adapting spiking (NASP), persistent stuttering (PSTUT), and persistent slow-wave bursting (PSWB).

**Figure 1.**
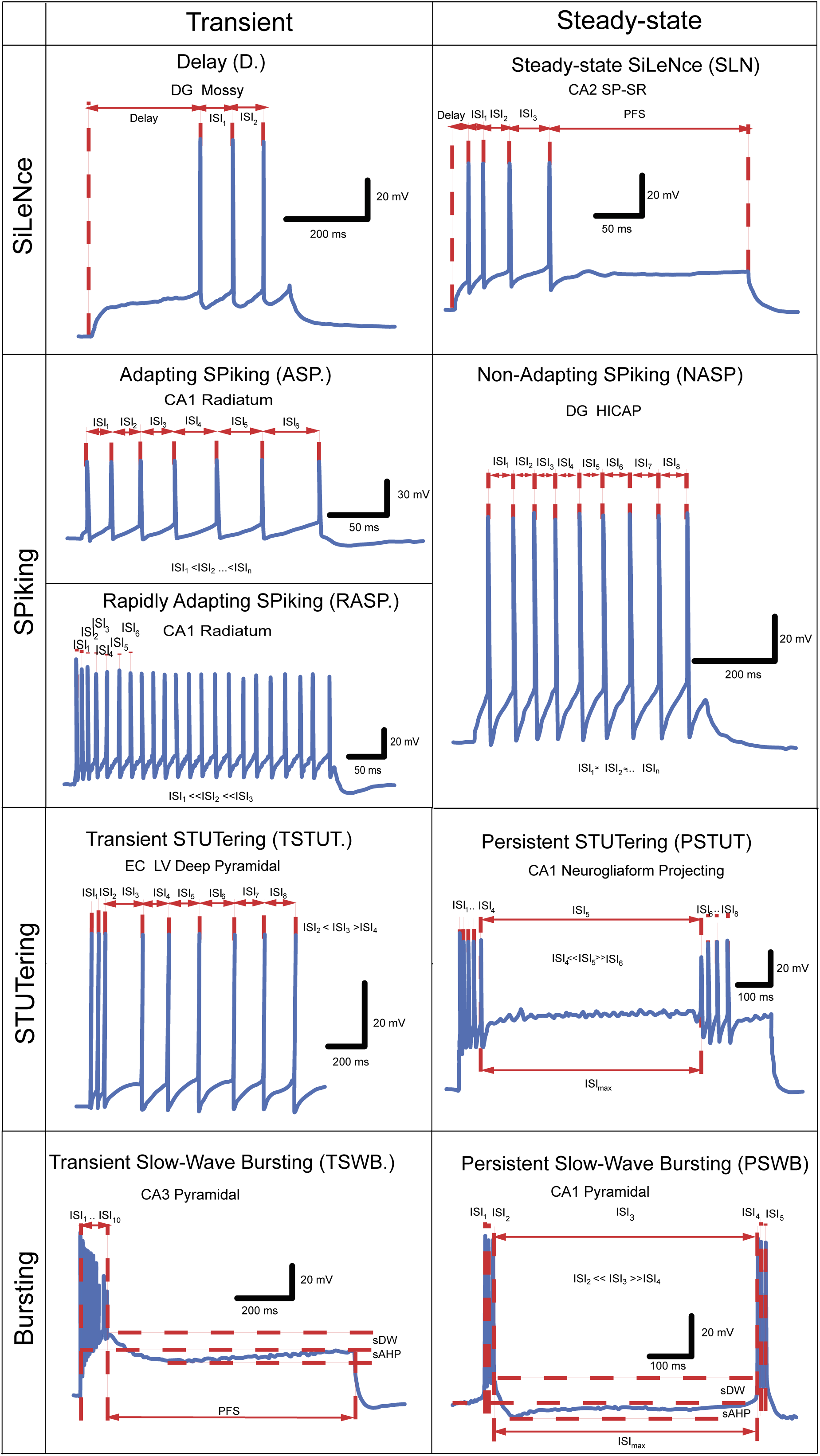
Firing pattern elements observable in hippocampal neurons *in vitro*. ISI - inter-spike interval, PFS – post firing silence, sDW – slow depolarization wave, sAHP – slow after-hyperpolarization. Original data extracted from Lübke et al. (1998) (D), Vida et al. (1998) (ASP), Pawelzik et al. (2002) (RASP), Hamam et al. (2002) (TSTUT), Chevaleyre and Seigelbaum (2010) (TSWB), Mercer et al. (2012) (SLN), Mott et al. (1997) (NASP), Fuentealba et al. (2010) (PSTUT), and Golomb et al. (2006) (PSWB, spontaneous bursting in Ca^2+^-free ACSF).

In certain cases, a constant current injection elicits firing patterns consisting of single firing pattern elements (NASP, PSTUT or PSWB). In other cases, complex firing patterns are observed as sequences of two or more firing pattern elements, such as delayed non-adapting spiking (D.NASP), silence preceded by adapting spiking (ASP.SLN), and non-adapting spiking preceded by delayed transient slow-wave bursting (D.TSWB.NASP). Experimental recordings without identifiable steady states were deemed uncompleted firing patterns (e.g. ASP., D.ASP., or RASP.ASP.).

In order to define the firing pattern elements unambiguously, we developed a set of quantitative classification criteria (Table 3). The transient response was classified as delayed (D.) if the latency to the first spike was longer than the sum of the first two inter-spike intervals (ISI_1_ and ISI_2_). Similarly, post-firing silence (PFS) was considered to be a steady state (SLN) if it exceeded the sum of the last two inter-spike intervals (ISI_n-1_ and ISI_n_). In addition, post-firing silence had to last at least twice the longest inter-spike interval (ISI_max_).

**Table 3.**
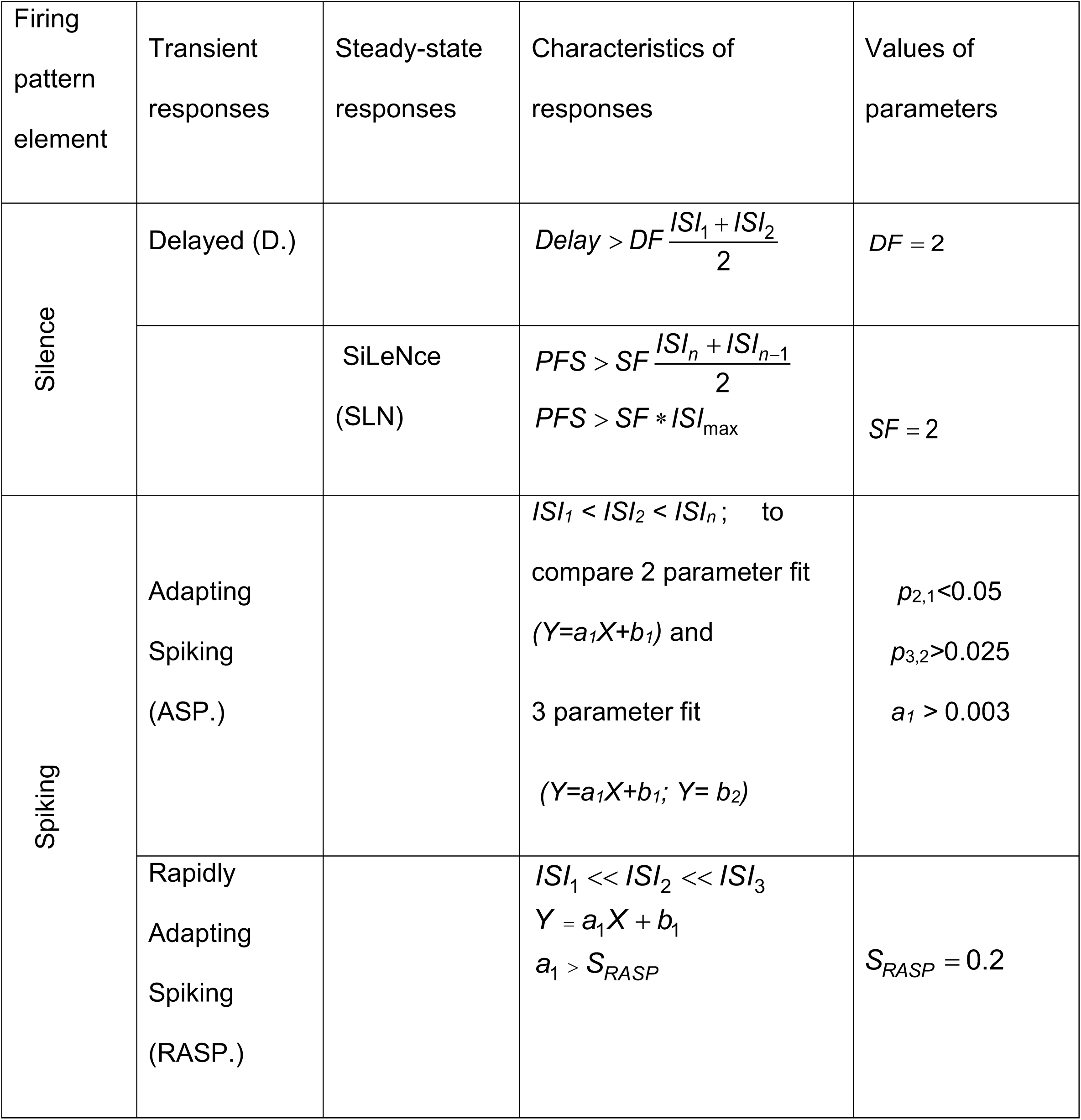
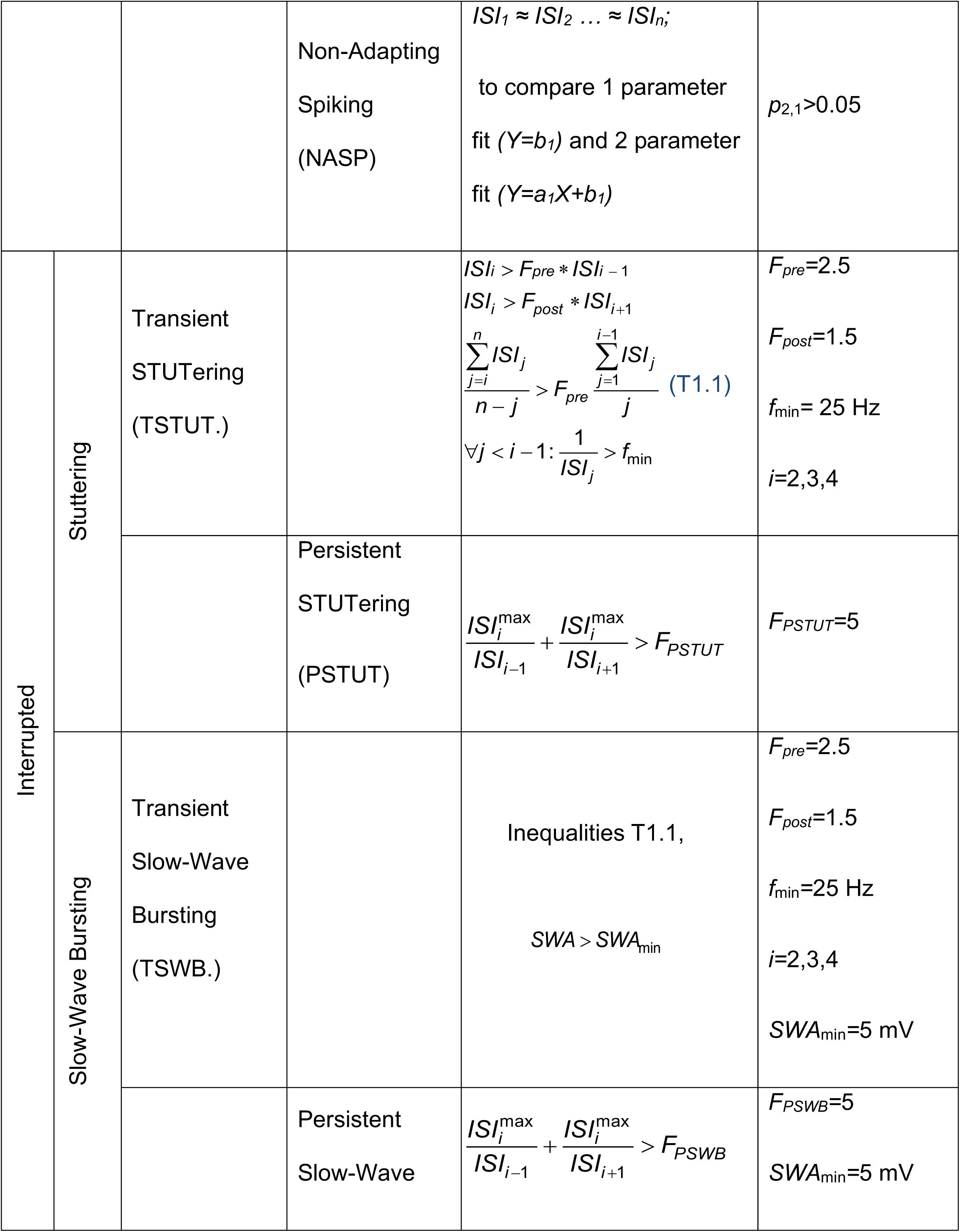
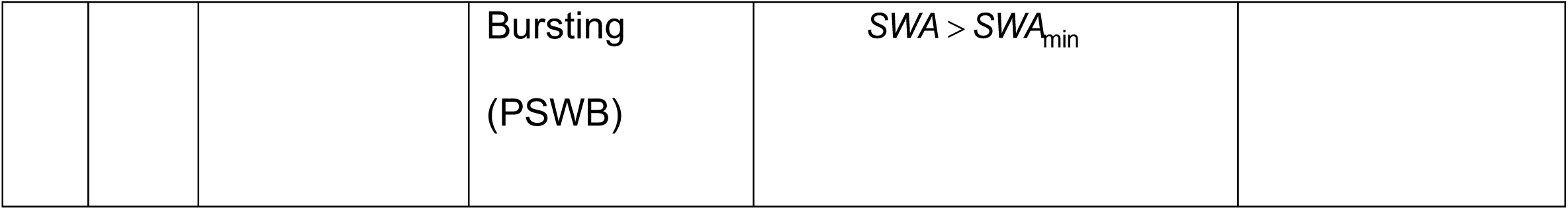
Principles of classification of firing pattern elements

| Firing pattern element | Transient responses | Steady-state responses | Characteristics of responses | Values of parameters |
| --- | --- | --- | --- | --- |
| Silence | Delayed (D.) | | $Delay > DF \frac{ISI_1 + ISI_2}{2}$ | $DF = 2$ |
| | | SiLeNce (SLN) | $PFS > SF \frac{ISI_n + ISI_{n-1}}{2}$<br>$PFS > SF * ISI_{max}$ | $SF = 2$ |
| Spiking | Adapting Spiking (ASP.) | | $ISI_1 < ISI_2 < ISI_n$; to compare 2 parameter fit<br>$(Y=a_1X+b_1)$ and<br>3 parameter fit<br>$(Y=a_1X+b_1; Y= b_2)$ | $p_{2,1} < 0.05$<br>$p_{3,2} > 0.025$<br>$a_1 > 0.003$ |
| | Rapidly Adapting Spiking (RASP.) | | $ISI_1 \ll ISI_2 \ll ISI_3$<br>$Y = a_1X + b_1$<br>$a_1 > S_{RASP}$ | $S_{RASP} = 0.2$ |

|  |  |  |  |  |  |
| --- | --- | --- | --- | --- | --- |
| | | | Non-Adapting Spiking (NASP) | $ISI_1 \approx ISI_2 \dots \approx ISI_n;$<br>to compare 1 parameter fit ($Y=b_1$) and 2 parameter fit ($Y=a_1X+b_1$) | $p_{2,1}>0.05$ |
| Interrupted | Stuttering | Transient STUTering (TSTUT.) | | $ISI_i > F_{pre} * ISI_{i-1}$<br>$ISI_i > F_{post} * ISI_{i+1}$<br>$\frac{\sum_{j=i}^n ISI_j}{n-j} > F_{pre} \frac{\sum_{j=1}^{i-1} ISI_j}{j}$ (T1.1)<br>$\forall j < i-1: \frac{1}{ISI_j} > f_{min}$ | $F_{pre}=2.5$<br>$F_{post}=1.5$<br>$f_{min}=25\text{ Hz}$<br>$i=2,3,4$ |
| | | | Persistent STUTering (PSTUT) | $\frac{ISI_i^{max}}{ISI_{i-1}} + \frac{ISI_i^{max}}{ISI_{i+1}} > F_{PSTUT}$ | $F_{PSTUT}=5$ |
| | Slow-Wave Bursting | Transient Slow-Wave Bursting (TSWB.) | | Inequalities T1.1,<br>$SWA > SWA_{min}$ | $F_{pre}=2.5$<br>$F_{post}=1.5$<br>$f_{min}=25\text{ Hz}$<br>$i=2,3,4$<br>$SWA_{min}=5\text{ mV}$ |
| | | | Persistent Slow-Wave | $\frac{ISI_i^{max}}{ISI_{i-1}} + \frac{ISI_i^{max}}{ISI_{i+1}} > F_{PSWB}$ | $F_{PSWB}=5$<br>$SWA_{min}=5\text{ mV}$ |
| | | | Bursting<br>(PSWB) | $SWA > SWA_{\min}$ | |

A persistent firing response with relatively equal inter-spike intervals denotes non-adapting spiking (NASP); in contrast, transients with a progressive increase or decrease of ISIs can be classified as adapting or accelerating spiking, respectively. To discriminate among several possible combinations of these firing patterns objectively and reproducibly, we devised a minimum information description criterion by comparing piecewise (segmented) linear regression models of increasing complexity. Specifically, non-adapting spiking (NASP) can be described by a single parameter, namely the (average) firing rate (*Y=c*). Similarly, fitting normalized inter-spike intervals versus normalized time with a (2-parameter) linear function *Y=aX+b* (with *a>0*) corresponds to adapting spiking (ASP.). Fitting data with a piecewise linear function

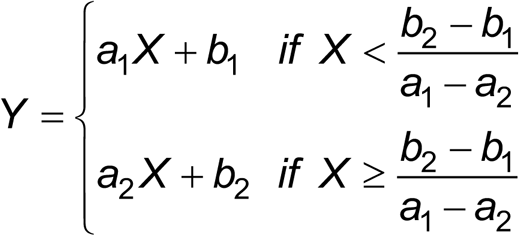

corresponds to adapting-non-adapting spiking (ASP.NASP) when *a*_*1*_*>0* and *a*_*2*_*=0* (3 parameters), and to adapting-adapting spiking with different adaptation rates (ASP.ASP.) when both *a*_*1*_*>0* and *a*_*2*_*>0* (4 parameters). We only selected a model with more parameters if the fit relative to a less complex model improved in a statistically significant way. The significance threshold for the differences between one-parameter fitting (NASP) and two-parameter linear-regression fitting (ASP.) was conventionally set at 0.05. Furthermore, in order to avoid identifying very weak adaptations as ASP., a minimum threshold of 0.003 was used for the slope *a*_*1*_.

For each subsequent stage of comparison, we used Bonferroni-corrected *p*-values. Specifically, in order for a pattern with an adapting spiking transient (i.e. ASP.) to be qualified as ASP.NASP, the *p*-value must be less than 0.025. Similarly, the *p*-value for the differences between three-parameter piecewise-linear-regression fitting (ASP.NASP) and four-parameter piecewise-linear-regression fitting (ASP.ASP.) must be less than 0.016. Figure 2 shows examples of fitting spiking activity with linear regression and piecewise linear regression models. If adaptation was only observed in the first two or three ISIs in a longer train of spikes, and if the linear fitting of slope *a*_*1*_ exceeded 0.2, then this transient was classified as rapidly adapting spiking (RASP.) (see Fig.1; cf. Pawelzik et al., 2002). For accelerating spiking (ACSP.), the linear fitting slope must be negative.

**Figure 2.**
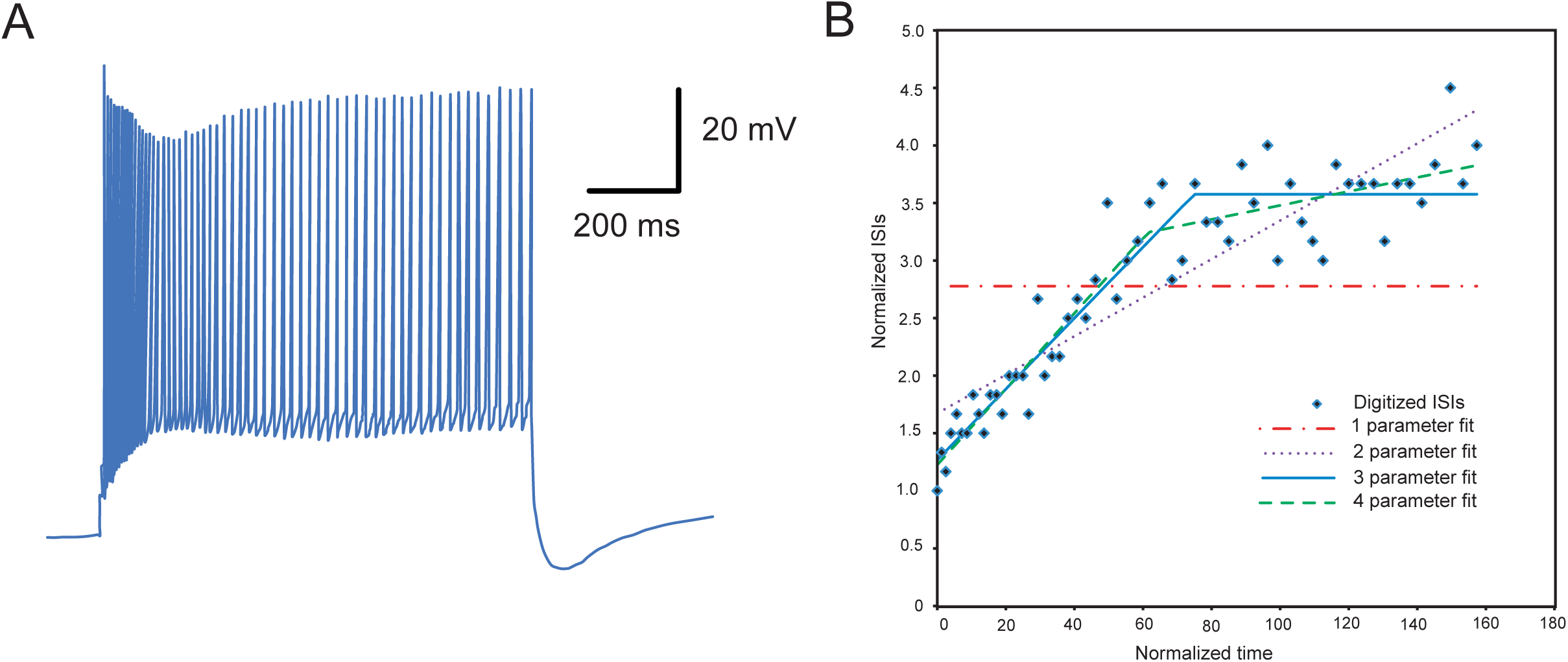
Examples of fitting of spiking activity with linear regression and piecewise linear regression models. **A.** Responses to current injection of a DG aspiny interneuron with axonal projection to the inner molecular layer (AIPRIM in Hippocampome.org) (Original data extracted from Lübke et al., 1998). **B.** Fitting of digitized experimental data with different models. 1 parameter fit is a constant function *Y=*2.78; 2 parameter fit is a linear function *Y=*0.017*X+1*.67; 3 parameter fit is a piecewise linear function 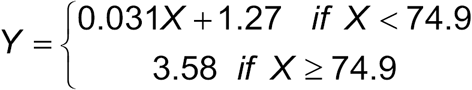; 4 parameter fit is a piecewise linear function 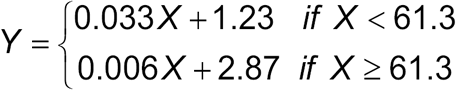. Based on *p*-values, the firing pattern was identified as adapting-non-adapting spiking (ASP.NASP): *p*_2,1_ < 0.05 (*p*_2,1_ =1.26·10^−10^), *p*_3,2_< 0.025 (*p*_3,2_=2.7·10^−3^), *p*_4,3_> 0.016 (*p*_4,3_=5.5·10^−2^). *p*_2,1_, *p*_3,2_, *p*_4,3_ – p-values of differences between 2 parameter fit and 1 parameter fit, 3 parameter fit and 2 parameter fit, 4 parameter fit and 3 parameter fit, respectively.

We defined transient stuttering (TSTUT.) as a short high-frequency (>25 Hz) cluster of action potentials (APs) followed by other distinctive activity. In addition, the first ISI after a TSTUT cluster must be 2.5 times longer than the last ISI of the cluster and 1.5 times longer than the next ISI (see Fig. 1; cf. Hamam et al., 2000). Under transient slow-wave bursting activity (TSWB.), a cluster of two or more spikes rides on a slow depolarization wave (>5mV) followed by a strong slow after-hyperpolarization (AHP) (see Fig. 1; cf. Chevaleyre and Siegelbaum 2010). Persistent stuttering (PSTUT) was classified as firing activity with high-frequency clusters of APs separated by silence intervals >5 times longer than the sum of the preceding and following ISIs (see Fig. 1; cf. Fuentealba et al. 2010; Price et al. 2005). Similarly, under persistent slow-wave bursting (PSWB) activity, these clusters of two or more tightly grouped spikes ride on slow depolarizing waves (>5 mV) followed by strong, slow AHPs (Golomb et al. 2006; Bilkey and Schwartzkroin 1990). As exemplified above, the choices of firing pattern identification parameters were consistent with literature reports of experimental results with similar activities.

### Algorithm Implementation

Based on the aforementioned methods, we implemented a firing pattern classification algorithm (Fig. 3) using the values of ISIs, delay, post-firing silence, and slow-wave amplitude as input data. Firing pattern elements were identified based on calculated characteristics of responses (Table 3). First, it was determined whether the pattern contained a delay (D.), then whether it contained a TSTUT. or TSWB. The remaining ISIs were processed using the described statistical test to identify spike frequency adaptation (ASP., ASP.NASP, ASP.ASP.) by fitting the sequence of intervals with a piecewise linear function. In the case of an incomplete pattern or an insufficient number of ISIs to perform the test, the presence of post firing silence (SLN) was checked. If the test did not identify the pattern containing the adaptation, then the firing pattern was checked for the presence of PSTUT or PSWB, and then for NASP or RASP. If rapid adaptation was detected, the cycle with the statistical test was performed again on the remaining ISIs. The algorithm terminated upon detection of one of the steady states (SLN, NASP, PSTUT, or PSWB).

**Figure 3.**
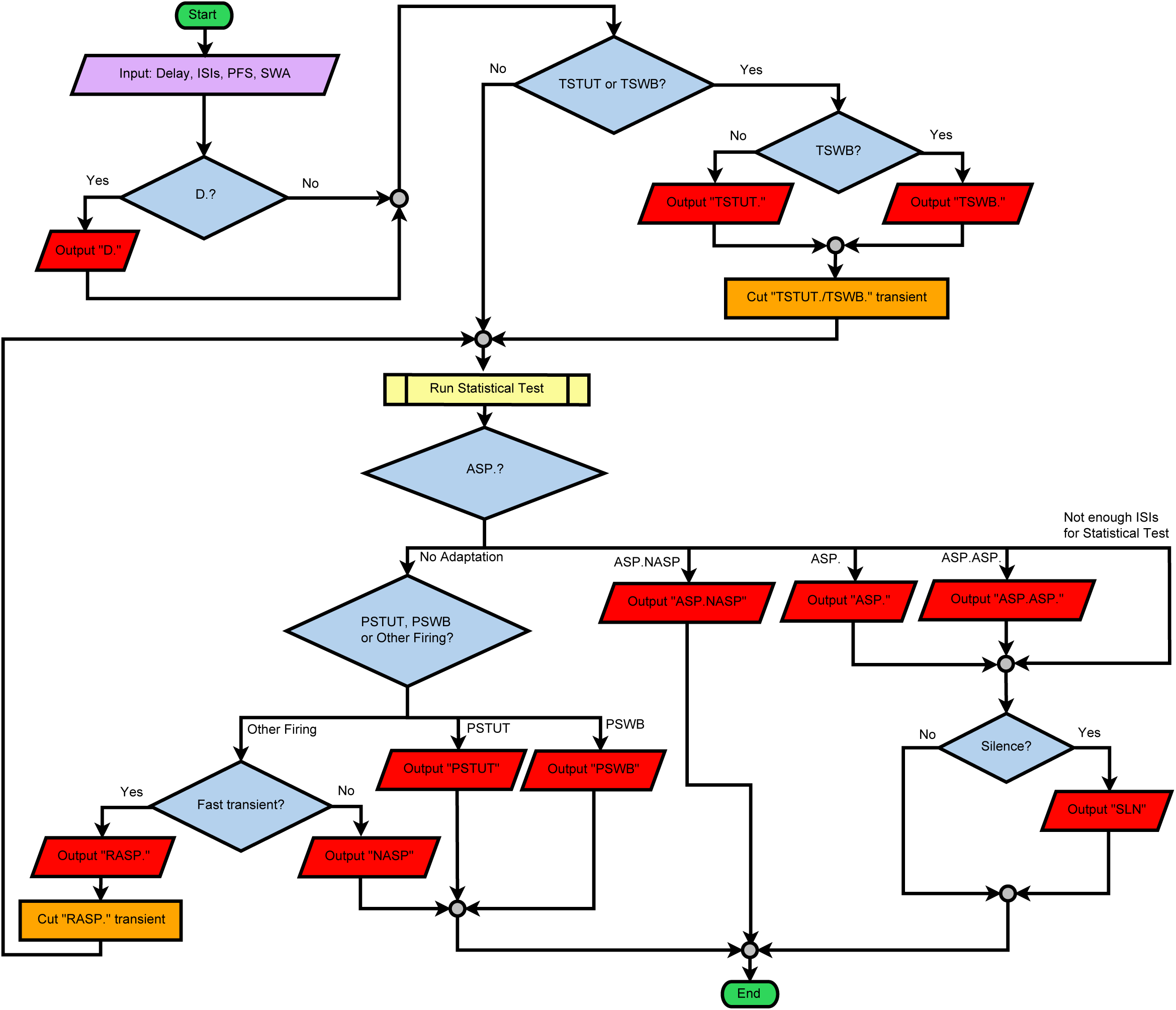
Flow chart of general procedure for firing pattern identification. See text for abbreviations. Source code and executable of Java implementation available at github.com/Hippocampome-Org/NeuroSpikePatterns.

### Software Accessibility

The classification algorithm was initially piloted in Microsoft Excel (Visual Basic) using Solver and the Data Analysis Toolbox (F-test and t-test) to perform piecewise linear fitting and statistical tests. The program was then re-implemented in the Java programming language using the Apache Commons Mathematics Library (commons.apache.org/proper/commons-math). The Java implementation is available open source at github.com/Hippocampome-Org/NeuroSpikePatterns.

### Experimental Design and Statistical Analysis

We explored pairwise correlations between all observed firing patterns, firing pattern elements, and 316 properties of Hippocampome.org neuron types, including: primary neurotransmitter; axonal, dendritic, and somatic locations in the 6 sub-regions and 26 parcels of the hippocampal formation; the projecting (between sub-regions) or local (within sub-regions) nature of axonal and dendritic patterns; axon and dendrite co-presence within any parcel; axonal and dendritic presence in a single layer only (intra-laminar) or in ≥3 layers (trans-laminar); clear positive or negative expression of any molecular markers; high (top third) or low (bottom third) values for biophysical properties (Wheeler et al., 2015); and potential connectivity patterns and super-patterns (Rees et al., 2016). To evaluate the correlations between these categorical properties, we used 2×2 contingency matrices with Barnard’s exact test (Barnard, 1947), which provides the greatest statistical power when row and column totals are free to vary (Lydersen et al., 2009). The correlation analysis was implemented in MATLAB (MathWorks, Inc.).

We analyzed numerical electrophysiological data, such as the relationship between the width of an action potential and the minimum ISI using linear regression and histograms. Spike duration was measured as the width at half-maximal amplitude as is most commonly defined (Bean, 2007). Minimum inter-spike intervals (ISI_min_) were extracted from digitized recordings or directly from tables or textual excerpts of the corresponding papers.

For cluster analysis of weighted categorical firing pattern data, we assigned weights to firing pattern elements according to the formula *W*_*e*_*=(N-n*_*e*_*)/N*, where *W*_*e*_ is the weight of the element *e, n*_*e*_ is the number of cell types expressing firing pattern(s) with element *e, N* is the total number of cell types/subtypes, and *e*={ASP., D., RASP., NASP, PSTUT, PSWB, SLN, TSUT., TSWB.}. We employed a two-step cluster analysis using the IBM SPSS Statistics 24 software. Silhouette measures of cohesion and separation greater than 0.5 indicated that the elements were well matched to their own clusters and poorly matched to neighboring clusters, and that the clustering configuration was appropriate.

Statistical data were expressed as mean ± standard deviation.

### Web portal and database representation of firing patterns and experimental conditions

Hippocampome.org provides access to morphological, molecular, electrophysiological, and connectivity information for 122 neuron types. The firing pattern data newly added and made freely available for download with this work include recording illustrations, the duration and amplitude of stimulation, digitized ISIs and firing pattern parameters (as comma-separated-value files), the complete solution compositions of the ACSF and of the micropipettes or patch pipettes, and the result of the firing pattern classification algorithm detailed above. Additional metadata is collected and displayed for all electrophysiological evidence in Hippocampome.org including the animal species (rat vs. mouse) and other details regarding the subject (inbred strain, age, sex, and weight, if reported), slice thickness and orientation, recording methods (intracellular microelectrode or variations of patch clamp), and temperature.

The implementation of Hippocampome.org supports the model-view-control software design. The model component defines the database interface and is provided solely by server-side code. The view component rendering the web pages and the control code implementing the decision logic are both served up by the server, but are run in the user’s browser. The underlying relational database ensures flexibility in establishing relations between data records.

Hippocampome.org is deployed on a CentOS 5.11 server running Apache 2.2.22 and runs on current versions of several web browsers (Mozilla Firefox, Google Chrome, Apple Safari, and Microsoft Internet Explorer). Knowledge base content is served up to the PHP 5.3.27 website from a MySQL 5.1.73 database. Django 1.7.1 and Python 3.4.2 provide database ingest capability of comma separated value annotation files derived from human-interpreted peer-reviewed literature. Hippocampome.org code is available at github.com/Hippocampome-Org.

## Results

### From firing patterns to firing pattern phenotypes

Version 1.3 of Hippocampome.org contains suitable electrophysiological recordings for 90 of the 122 morphologically identified neuron types. Applying the firing pattern identification algorithm to these digitized data resulted in the detection of 23 different firing patterns. A given neuron type may demonstrate distinct firing patterns in response to different stimuli or conditions. The set of firing patterns exhibited by a given neuron type forms its firing pattern phenotype.

The simplest case consists of those neuron types that systematically demonstrate the same firing pattern independent of experimental conditions or stimulation intensity. These neuron types may still display quantitatively different responses to stimuli of various amplitudes (typically increasing their firing frequency upon increasing stimulation), but their qualitative firing patterns remain the same. We identified 37 such “individual/simple-behavior types” in Hippocampome.org, as exemplified by DG Basket cells with their NASP phenotype (Savanthrapadian et al., 2014).

In contrast to the above scenario, certain neuron types exhibit qualitatively distinct firing patterns in response to different amplitudes of stimulation. We identified 21 such “multi-behavior” types; for instance, CA1 Neurogliaform cells (Price et al., 2005; Tricoire et al., 2011) display delayed firing, adapting spiking, and persistent stuttering at different stimulus intensities. The firing phenotype of these interneurons thus consists of the combination of all three firing patterns.

In a different set of cases, subsets of neurons from the same morphologically identified type display distinct firing patterns under the same experimental conditions (typically from the same study) in response to identical stimulation. These neuron types can thus be divided into electrophysiological subtypes. For example, of the CA3 Spiny Lucidum interneurons, some are adapting spikers whereas others are persistent stutterers (Szabadics and Soltesz, 2009). In certain neuron types, one or more of the subtypes could also display multiple behaviors at different stimulation intensities. For instance, a subset of entorhinal Layer III Pyramidal neurons consists of non-adapting spikers and another subset switches from ASP.NASP at rheobase to RASP.ASP. at higher stimuli (Canto and Witter, 2012b). Of the 90 neuron types with firing patterns in Hippocampome.org, 22 could be divided into 52 electrophysiological subtypes. Notably, these included the principal neurons of most sub-regions of the hippocampal formation: CA3, CA1, and subiculum Pyramidal cells, entorhinal Spiny Stellate cells, but also several GABAergic interneurons such as Total Molecular Layer (TML) cells (Mott et al, 1997). Specifically, 8 neuron types yielded 18 subtypes exclusively demonstrating single behaviors; for 11 neuron types, at least one of the subtypes exhibited multi-behaviors, resulting in 13 multi-behavior subtypes and 13 additional single-behavior subtypes.

This meta-analysis is complicated by the variety of experimental conditions used in the published literature from which the electrophysiological data were extracted. Several differences in materials and methods could affect firing patterns above and beyond common species (rats vs. mice) or recording (patch clamp vs. microelectrode). For example, 30% of experimental traces were recorded from transverse slices, 24% from horizontal, 8% coronal, 29% mixed (e.g. “horizontal or semicoronal”), and 9% other directions (e.g. custom angles). Furthermore, pipettes were filled with potassium gluconate in 69% of cases, with potassium methylsulfate in 22%, and with potassium acetate in 9% (see e.g. Table 2). While these different experimental conditions can affect membrane biophysics substantially (Tebaykin et al., 2018) and often quantitatively influence neuronal firing (e.g. changing the spiking frequency), occasionally they can also cause a qualitative switch between distinct firing patterns. A striking case is that of rat DG Granule cells, which have demonstrated transient slow-wave burst followed by silence in whole-cell recordings of horizontal slices from Sprague-Dawley animals (Williams et al., 2007); delayed non-adapting spiking in whole-cell recording of transverse slices from Wistar animals (Lübke et al., 1998); or adapting spiking in intracellular recording of horizontal slices from Wistar animals (Han et al., 1993). Because the different firing patterns could be caused by the differences in experimental methods, we annotate a possible “condition-dependence,” but cannot conclusively categorize these cells as multi-behavior or subtypes. Most of the condition-dependent behaviors could be attributed at least in part to the occasional use of microelectrode instead of patch-clamp (now considered the preferred recording method) or the animal species as in the case of CA1 Horizontal Basket cells which display adapting and non-adapting firing in rats and mice, respectively (Zemankovics et al, 2010; Tricoire et al, 2011).

Condition dependence can alter the firing patterns not only in cell types with single behaviors, such as MOPP cells (Han et al., 1993; Armstrong et al., 2011), but also in multi-behavior neuron types, such as CA1 Axo-Axonic cells (Buhl et al., 1994; Pawelzik et al., 2002). These cases account for 6 and 4 Hippocampome.org neuron types, respectively. Lastly, condition dependence may also be found in specific electrophysiological subtypes, whether they display single behaviors, such as CA1 Pyramidal neurons (Chevaleyre and Siegelbaum, 2010; Zemankovics et al, 2010; Kirson and Yaari, 2000; Staff et al., 2000) or multi-behavior, such as entorhinal Layer V Deep Pyramidal neurons (Canto and Witter, 2012; Hamam et al., 2000; Hamam et al., 2002). These cases respectively account for 2 and 1 Hippocampome.org neuron types, in turn giving rise to 6 condition-dependent subtypes with single behaviors and 2 condition-dependent subtypes with multi-behavior.

Figure 4 presents the full firing pattern phenotypes of all 90 Hippocampome.org neurons with available data in form of separate matrices for the 68 individual neuron types (Fig. 4A) and the 52 subtypes divided from the remaining 22 types (Fig. 4B). In both cases the simple behaviors constitute larger proportions than multi-behavior with condition dependence only reported for a minority of types and subtypes (Fig. 4C). Across these neuron types/subtypes, 44 distinct phenotypes can be identified as unique combinations of firing patterns, excluding those that differ from others only by the absence of a detectable stable state in one of the firing patterns (like ASP. versus ASP.NASP or ASP.SLN). An interactive online version of these matrices is available at hippocampome.org/firing_patterns.

**Figure 4.**
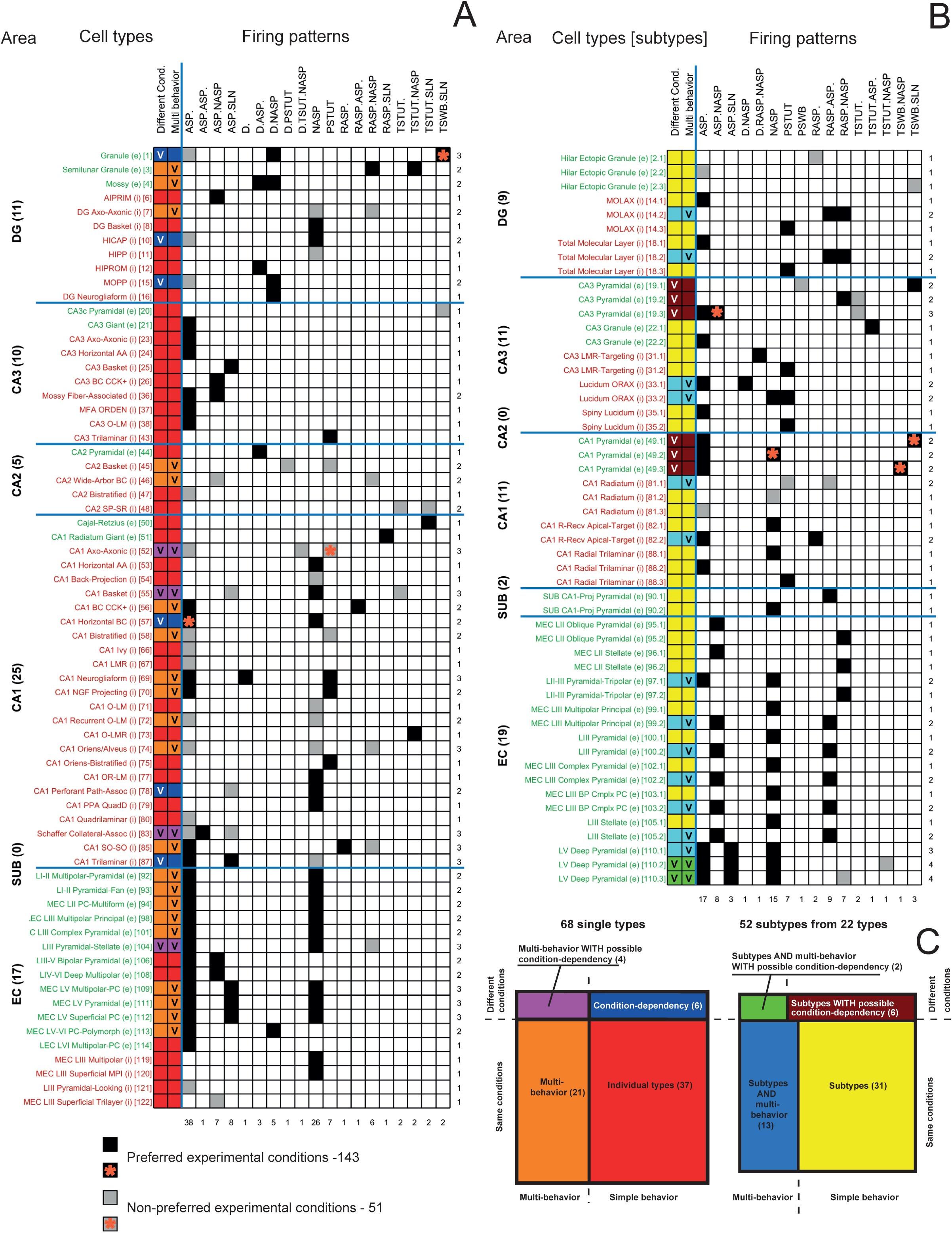
Identified firing patterns and firing pattern phenotypes complexity of neuron types (**A**) and subtypes (**B**). Online matrix: hippocampome.org/firing_patterns. Green and red cell type/subtype names denote excitatory (e) and inhibitory (i) neurons, respectively. FPP is firing pattern phenotype. The numbers in the brackets correspond to the order in which the cell types were presented in the Hippocampome (ver. 1.3). The orange asterisk denotes different experimental conditions. **C**. Complexity of firing pattern phenotypes; percentages and ratios indicate occurrences of phenotypes of different complexity among 120 cell types/subtypes.

### Dissecting firing patterns into firing pattern elements across neuron types

Firing patterns and firing pattern elements are also diverse with respect to their relative frequency of occurrence among hippocampal neuron types. Firing patterns can be grouped based on the number of elements comprising them, namely single (e.g., NASP or PSTUT), double (e.g. ASP.NASP or TSWB.SLN), and triple (D.RASP.NASP and D.TSWB.NASP) or based on whether they are completed (ASP.NASP, TSWB.SLN) or uncompleted, as in ASP., RASP.ASP., and TSTUT.ASP. (Fig. 5A). Of the nine firing pattern elements, the most frequent are ASP and NASP, while the least common are TSTUT, TSWB, and PSWB (Fig. 5B). Notably, accelerated spiking (ACSP) has not been reported in the rodent hippocampus although it is commonly observed in other neural systems, such as turtle ventral horn interneurons (Smith and Perrier 2006) and motoneurons (Leroy et al. 2014).

**Figure 5.**
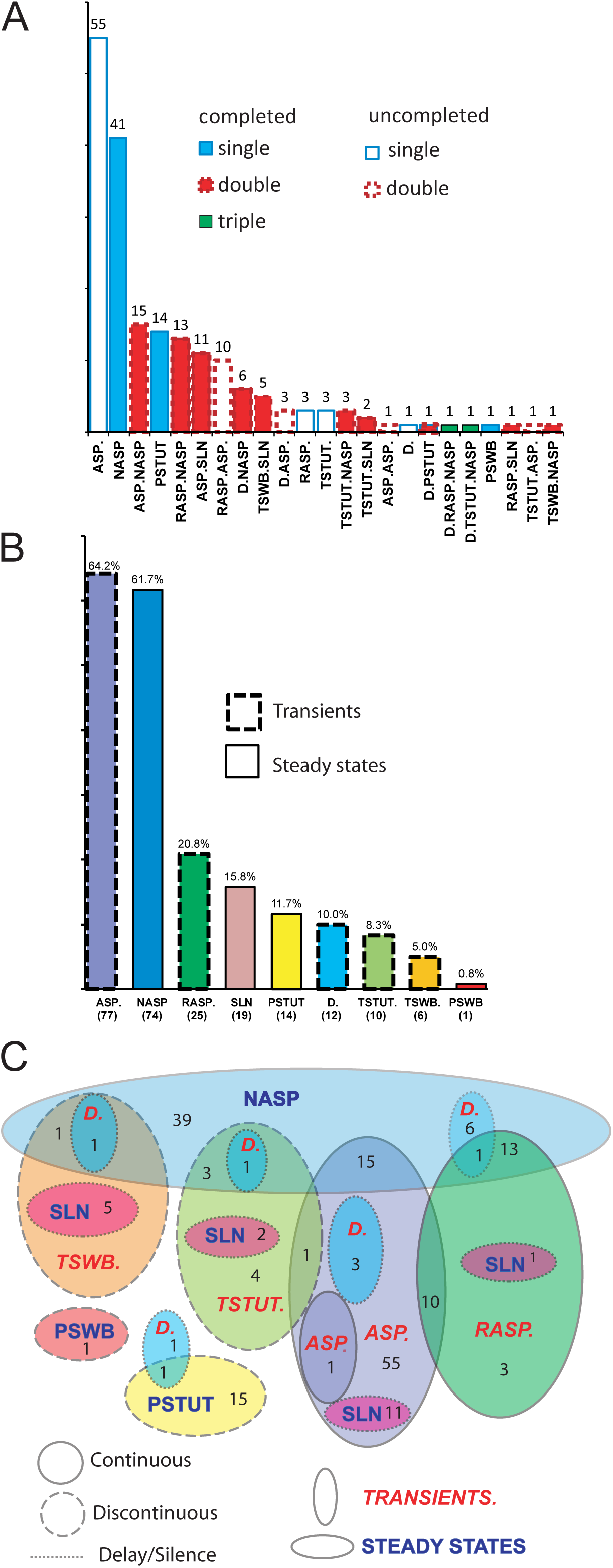
Occurrence of firing patterns, firing pattern elements and firing pattern phenotypes among the hippocampal formation neuron types. **A**. Distribution of 23 firing patterns; total numbers are shown above the bars. **B**. Distribution of 9 firing pattern elements; total numbers are in parentheses below and percentages of occurrence among 90 cell types are above the bars. **C**. Relationships between firing pattern elements in the firing patterns of hippocampal neuron types. Numbers of cell types with distinctive firing patterns are indicated.

The relationships between sets of firing pattern elements observed in hippocampal neuron types can be summarized in a Venn diagram with firing pattern elements represented as ellipses and the intersections thereof corresponding to complex firing patterns (Fig. 5C). This analysis highlights the following features: the four main firing transients (ASP., RASP., TSTUT., TSWB.) often end either with NASP or with SLN; ASP. is often preceded by RASP. and occasionally by TSTUT.; interrupted steady-state firings (PSTUT and PSWB) stand out as a separate group; and delay (D.) most often precedes NASP.

Our classification of firing pattern elements implies the possibility of three completed single-element firing patterns (NASP, PSTUT, PSWB) and 19 completed double-element firing patterns consisting of one of four steady states (SLN, NASP, PSTUT, PSWB) preceded by one of five transients (D, ASP, RASP, TSTUT, TSWB), with exclusion of the “empty” combination D.SLN. Also, four double-transients are possible after an initial delay, resulting in an additional 16 triple-element firing patterns. Only 15 of these possible 38 completed firing patterns were discovered in literature data for morphologically identified hippocampal neuron types (Table 4). Three additional firing patterns were found in other neurons: D.PSWB has been shown in the cultured *rutabaga* mutant giant neuron of *Drosophila* (Zhao and Wu 1997), D.ASP.SLN in the neuron of the external lateral subnucleus of the lateral parabrachial nucleus (Hayward and Felder 1999), and D.TSTUT.SLN in the striatal fast-spiking neuron (Sciamanna and Wilson 2011). We deemed 16 firing patterns as “not found but possible” (white shading and black text in Table 4) and 4 firing patterns as “improbable” (white shading and gray text). In particular, we consider combination of stuttering and slow-wave bursting (TSWB.PSTUT or TSTUT.PSWB) as unlikely to occur under physiological conditions from a dynamical viewpoint due to incompatible underlying mechanisms. Slow-wave bursting is provided by a slow negative feedback which terminates the burst of action potential evoking slow AHP. Such feedback could be produced by different ionic mechanisms, but it is most typically based on intracellular Ca^2+^ dynamics and Ca^2+^-activated K^+^ current (Golomb et al. 2006; Xu and Clancy 2008) or muscarinic-sensitive K^+^ current (Golomb et al. 2006). Slow-wave bursting could be “square-wave bursting”, with one slow process, or “parabolic bursting”, with two (positive and negative feedback) slow processes (Rinzel and Ermentrout 1998). In contrast, stuttering activity is associated with “elliptic bursting” (Golomb et al. 2007), where the silent phase is characterized by dumping and growing fast (spiking) oscillations as the trajectory slowly drift through bifurcation of the fast subsystem (Rinzel and Ermentrout 1998). Suggested mechanism for stuttering in fast spiking interneurons includes Na^+^ “window” current that induces high frequency tonic firing, and slowly inactivating K^+^ current through KV1 channels (Golomb et al. 2007).

**Table 4.**
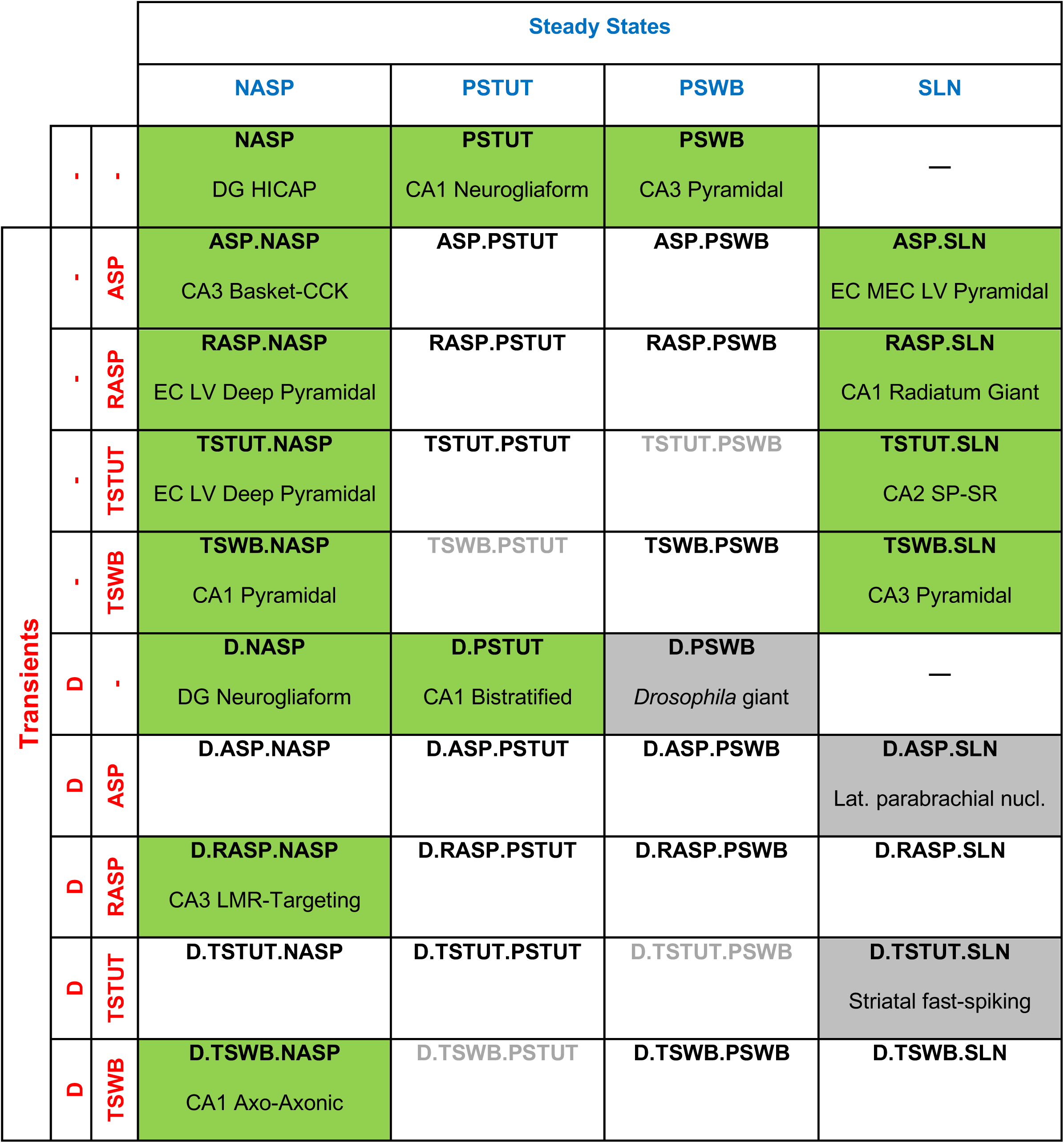
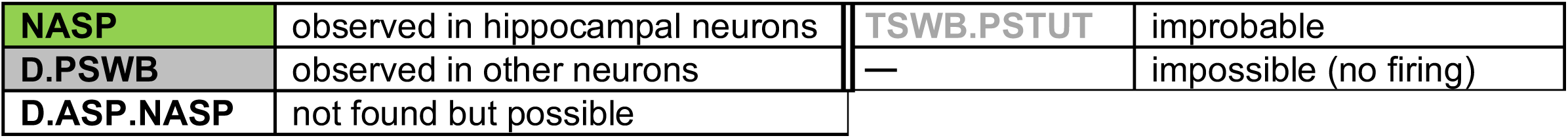
Occurrences of completed firing patterns in hippocampal and other neurons

### Classification and distribution of firing pattern phenotypes

In order to classify the 44 unique firing pattern phenotypes observed in the hippocampal formation, we weighted the constituent firing pattern elements according to the frequency of occurrence among 120 neuron types and electrophysiological subtypes (see *Materials and Methods*). As a result, infrequent firing pattern elements (PSWB, TSTUT and TSWB) received high weights (0.99, 0.95 and 0.93, respectively), very frequent elements (ASP and NASP) were assigned low weights (0.42 and 0.41), and common elements (D, RASP, PSTUT and SLN) obtained intermediate weights (0.90, 0.80, 0.88 and 0.87). Two-step cluster analysis identified ten firing pattern families as leaves of a seven-level hierarchical binary tree (Fig. 6A). At the highest level, hippocampal neuron types and subtypes are divided into two major groups: those with spiking phenotypes (78%) and those with interrupted firing phenotypes (22%). The latter are separated into bursting (6%) and stuttering (16%), and each of these is subdivided into persistent and non-persistent families. A first group of the neuron types with spiking phenotypes is distinguished based on delay (9% of cell types). The remaining neuron types split into adapting (54%) and non-adapting phenotypes (15%). The adapting group consists of neuron types with rapidly adapting phenotypes (18%) and normally adapting (36%) phenotypes. Among the normally adapting group, the following phenotypes can be distinguished: discontinuous adapting spiking (6%) with ASP.SLN pattern, adapting-non-adapting spiking (15%) with ASP.NASP patterns, and a last “spurious” phenotype of uncompleted adapting spiking (15%) with ASP. pattern only, for which the steady state (SLN or NASP) was not determined. This division of the adapting spiking groups reflects differences in adaptation rates, duration, and subsequent steady states.

**Figure 6.**
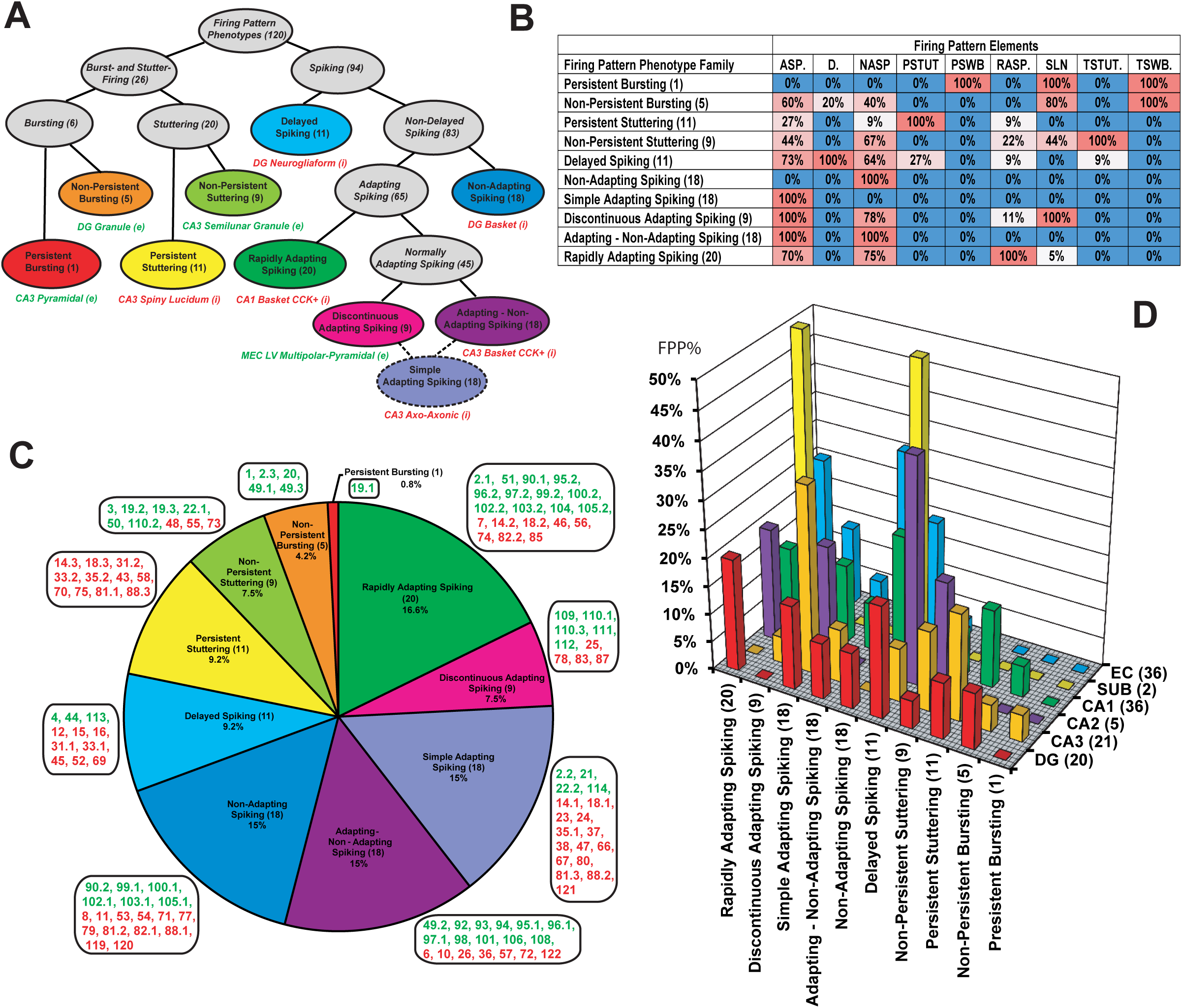
Ten firing pattern phenotype families from 120 neuron types/subtypes. **A**. Hierarchical tree resulting from two-step clustering of weighted firing pattern elements with representative examples of cell types/subtypes which belong to corresponding firing pattern phenotype family. Simple adapting firing pattern phenotype is not unique (see *Results*). **B**. Percentage of occurrence of firing pattern elements in families of firing pattern phenotypes. **C**. Relative proportions of firing pattern phenotypes among neuron types/subtypes. Green and red numbers represent excitatory and inhibitory cell types/subtypes as enumerated in Fig. 4. **D**. Distribution of firing pattern phenotypes in sub-regions of the hippocampal formation. FPP% is percentage of expression of firing pattern phenotypes.

This analysis also highlights the most distinguishing firing pattern elements of each family (Fig. 6B). In particular, D. is the defining element for delayed spiking, PSTUT for persistent stuttering, ASP. and SLN for discontinuous adapting spiking. Each of the four major elements of interrupted firing patterns (PSWB, PSTUT, TSWB. and TSTUT.) is observed in a single firing pattern phenotype (persistent bursting, non-persistent bursting, persistent stuttering, and non-persistent stuttering, respectively). Other firing pattern elements (D., RASP., ASP., NASP, and SLN) appear in several firing pattern phenotypes. The proportions of non-defining firing pattern elements range from 5% to 83%.

The families of firing pattern phenotypes are differentially distributed within the set of 120 neuron types/subtypes (Fig. 6C). Certain phenotype families are associated with excitatory neuron types, either exclusively (e.g. persistent bursting and non-persistent bursting) or predominantly (non-persistent stuttering, rapidly adapting, and adapting-non-adapting spiking). Conversely, persistent stuttering, delayed spiking, non-adapting spiking and simple adapting spiking are phenotypes composed largely by inhibitory neuron types. The discontinuous adapting spiking phenotype has relatively balanced proportions of excitatory and inhibitory neuron types.

The firing pattern phenotypes also have different distributions among the sub-regions of the hippocampal formation (Fig. 6D). Among CA1 neuron types, the persistent stuttering (16%), non-adapting (24%), simple adapting (16%), and rapidly adapting spiking (13%) phenotypes are more common than other phenotypes; in DG, the most expressed phenotypes are delayed (20%), rapidly adapting (20%), and simple adapting spiking (15%); in EC, ASP-NASP (61%), discontinuous ASP. (11%), RASP. (28%), and NASP (19%) occur more often than other phenotypes.

### Usage of information from Hippocampome.org

#### Searching and Browsing

The addition of firing pattern data to Hippocampome.org extends opportunities for broad-scope analytics and quick-use checks of neuron types. Similar to morphological, molecular, and biophysical information, firing patterns and their parameters can be browsed online with the interactive versions of the matrices presented in Figure 4 (hippocampome.org/firing_patterns), along with an accompanying matrix to browse the stimulation parameters (duration and intensity) and the firing pattern parameters (delay, number of inter-spike intervals, etc.).

Moreover, all classification and analysis results reported here can be searched with queries containing AND & OR Boolean logic using an intuitive graphical user interface (see Hippocampome.org → Search → Neuron Type). The integration within the existing comprehensive knowledge base enables any combination of both qualitative (e.g. PSTUT) and quantitative (e.g. 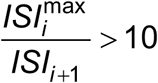) firing pattern properties with molecular (e.g. calbindin-negative), morphological (e.g. axons in CA1 pyramidal layer), and biophysical (e.g. action potential width < 1 ms) filters (Fig. 7). For example, of 14 neuron types with persistent stuttering, in 5 the maximum inter-spike interval is at least an order of magnitude longer than the subsequent spike. When adding the other three selected criteria, the compound search leads to a single hit: CA1 Axo-axonic neurons (Fig. 7A). Clicking on this result leads to the interactive neuron page (Fig. 7B) where all information associated with a given neuron type is logically organized, including synonyms, morphology, biophysical parameters, molecular markers, synaptic connectivity, and firing patterns. Every property on the neuron pages and browse matrices, including firing patterns and their parameters, links to a specific evidence page that lists all supporting bibliographic citations, complete with extracted quotes and figures (Fig. 7C). The evidence page also contains a table with all corresponding firing pattern parameters (Fig. 7D), experimental details including information about animals (Fig. 7E), preparations (Fig. 7F), recording method and intra-pipette solution (Fig. 7G), ACSF (Fig. 7H), and a downloadable file of inter-spike intervals (Fig. 7I).

**Figure 7.**
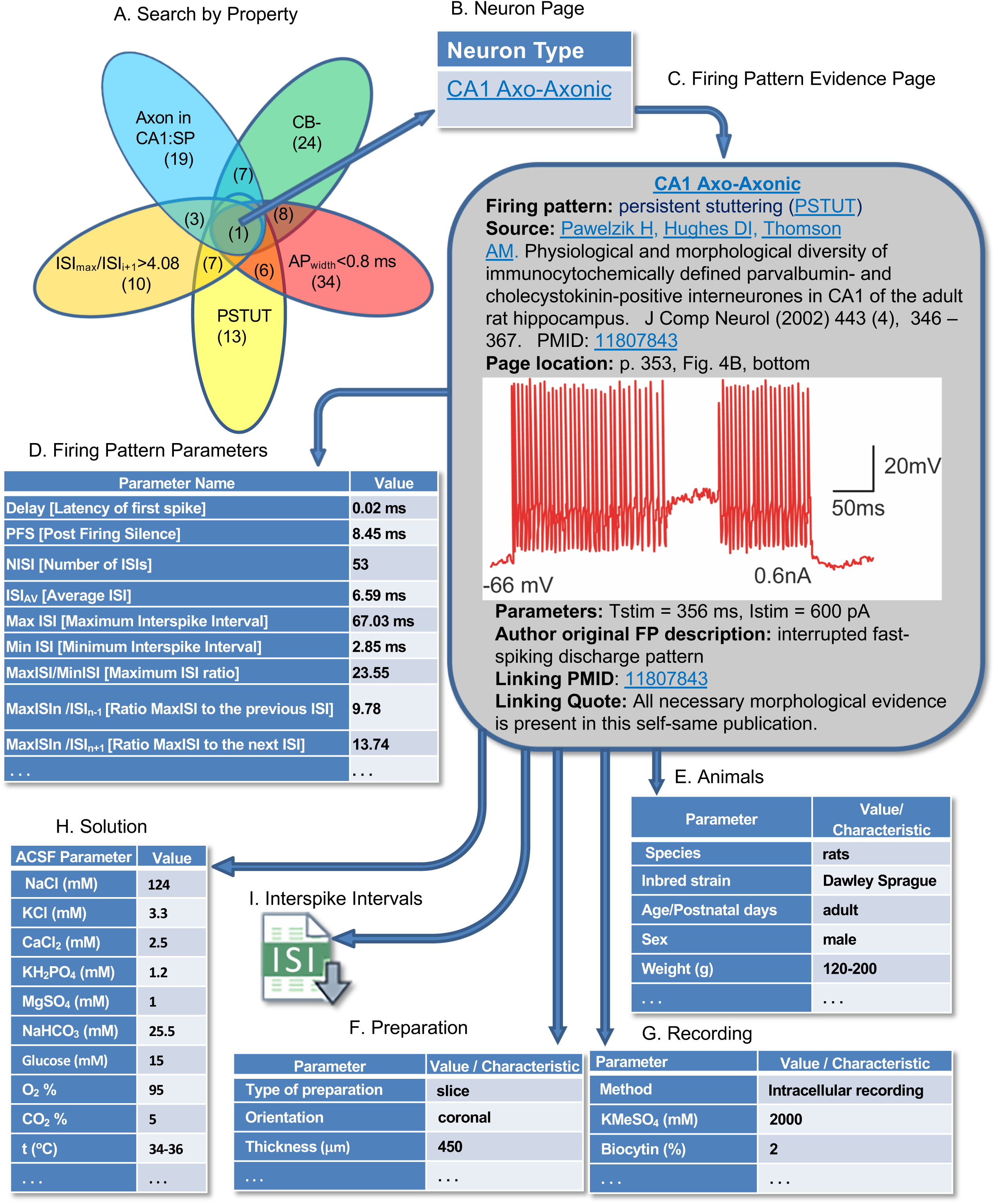
Hippocampome.org enables searching neuron types by neurotransmitter; axon, dendrite, and soma locations; molecular expression; electrophysiological parameters; input/output connectivity; firing patterns, and firing pattern parameters. **A**. Sample query for calbindin-negative neuron types with axons in CA1 stratum pyramidale, AP_width_ <0.8 ms, PSTUT firing, and ratio of maximum ISI to the next ISI greater than 4.8. Numbers in parentheses indicate the number of neuron types with the selected property or specific combination of properties. **B**. Search results are linked to the neuron page(s). **C**. The neuron page is linked to the firing pattern evidence page. Original data extracted from Pawelzik et al., 2002. **D-H.** All firing patterns parameters (**D**), experimental details including information about animals (**E**), preparations (**F**), recording method and intra-pipette solution (**G**), as well as ACSF composition (**H**) can be displayed. **I.** Downloadable comma-separated-value file of inter-spike intervals.

The portal also reports, when available, the original firing pattern name descriptions used by the authors of the referenced publication (Hippocampome.org → Search → Original Firing Pattern).

#### Statistical analysis of categorical data

Firing pattern information more than doubles the Hippocampome.org knowledge base capacity to over 27,000 pieces of knowledge, that is, associations between neuron types and their properties. This extension allows the unearthing of hidden relationships between firing patterns and molecular, biophysical, and morphological data in hippocampal neurons, which are otherwise difficult to find amongst the large body of literature. Several interesting examples of such findings are presented in Box 1. For instance, adapting spiking (ASP.) tends to co-occur with expression of cholecystokinin (p=0.0113 with Barnard’s exact test from all n = 26 pieces of evidence); moreover among 284 analyzed recordings there are no neurons with extremely high input resistance that show persistent stuttering (PSTUT) (p=0.0478).

**Box 1.** Examples of statistically significant correlations between 1196 firing patterns and 1197 known molecular, morphological and electrophysiological properties in 1198 hippocampal neurons

1. None of the 35 **glutamatergic** neuron types show persistent stuttering (**PSTUT**) (p=0.0083). Moreover, none of the neurons with high input resistance (**Rin**) display this steady state (p=0.0478). Thus, all PSTUT cells are GABAergic interneurons with low or intermediate input resistance.
2. Neither any of the 63 **non-projecting** (local circuit) neurons nor any of the 55 **GABAergic** neuron types display transient slow-wave bursting (**TSWB**.) (p=0.0214 and p=0.0215, respectively). Moreover, no neuron type or subtype is found with both TSWB firing and high values of hyperpolarization-induced **sag** potential (p=0.035). In other words, TSWB cells in the hippocampus are a subset of projecting (long-range) glutamatergic neurons with medium or low sags.
3. None of the 15 neuron types that express neuropeptide Y (**NPY**) become silent (**SLN**) after short firing discharge (p=0.0037). In contrast, half of the 14 NPYnegative cells demonstrate this steady state.
4. All of the 10 neuron types that express cholecystokinin (**CCK**) and the overwhelming majority of neuron types with high input resistance (17/18) display adapting spiking (**ASP.**) (p=0.0113). In contrast, this transient state is observed in just above half of CCK-negative cells (9/16) and two-thirds of cells with low or intermediate **R_in_** (12/18).
5. Of 14 neuron type with **wide AP**, only one (EC LV-VI Pyramidal-Polymorphic) shows delayed (**D.**) firing (p=0.021). In contrast, nearly half of neuron types without wide AP demonstrate this transient state.
6. With the exceptions of DG Semilunar Granule and CA1 O-LMR, none of the neurons with high threshold potential (**V_thresh_**) display transient stuttering (**TSTUT.**) (p=0.0481); similarly, none of the neurons with high amplitude of fast afterhyperpolarization (**fAHP**), except CA1 Cajal-Retzius, demonstrate TSTUT. (p=0.0098).

The p values are computed using Bernard’s exact test for 2 × 2 contingency tables (see *Materials and Methods*).

#### Analysis of numerical electrophysiological data

The extracted quantitative data allow one to study the relationship between firing pattern parameters and membrane biophysics or spike characteristics, such as the correlations between minimum inter-spike intervals (ISI_min_) and action potential width (AP_width_). We analyzed these two variables in the 81 neuron types and subtypes for which both measurements are available (Fig. 8). The scatter plot of AP_width_ against ISI_min_ reveals several distinct groupings (Fig. 8A), and the corresponding histograms (Figs. 8B,C) demonstrate poly-modal distributions. The horizontal dashed line (ISI_min_=34 ms) separates 9 neurons with slow spikes (all excitatory except one) from 72 neurons (61% of which are inhibitory) with fast and moderate spikes. The latter group shows a general trend of ISI_min_ rise with increasing AP_width_ (black dashed line in panel A). This trend was adequately fit with a linear function Y = 13.79X −0.05 (R = 0.72; p=0.03). Neuron types with slow spikes demonstrate the opposite trend, which was fit with a decreasing linear function Y = - 26.72X + 76.42 (R = −0.91, p=10^−6^). Furthermore, the neuron types can be separated by spike width. The vertical dashed lines *w1* (AP_width_=0.73 ms) and *w2* (AP_width_=1.12 ms) separate neuron types with narrow, medium and wide action potentials. The group of neuron types with narrow spikes (n=22) includes only inhibitory neurons, which have AP_width_ in the range from 0.20 to 0.73 ms (0.54 ± 0.12 ms). In contrast, the group of neuron types with wide spikes (n=28) contains only excitatory neurons with AP_width_ in the range from 1.13 to 2.10 ms (1.49 ± 0.23 ms). The group of neuron types with medium spikes (n = 31), with AP_width_ range from 0.74 to 1.12 ms (0.89 ± 0.12 ms), includes a mix of inhibitory (74%) and excitatory (26%) neurons.

**Figure 8.**
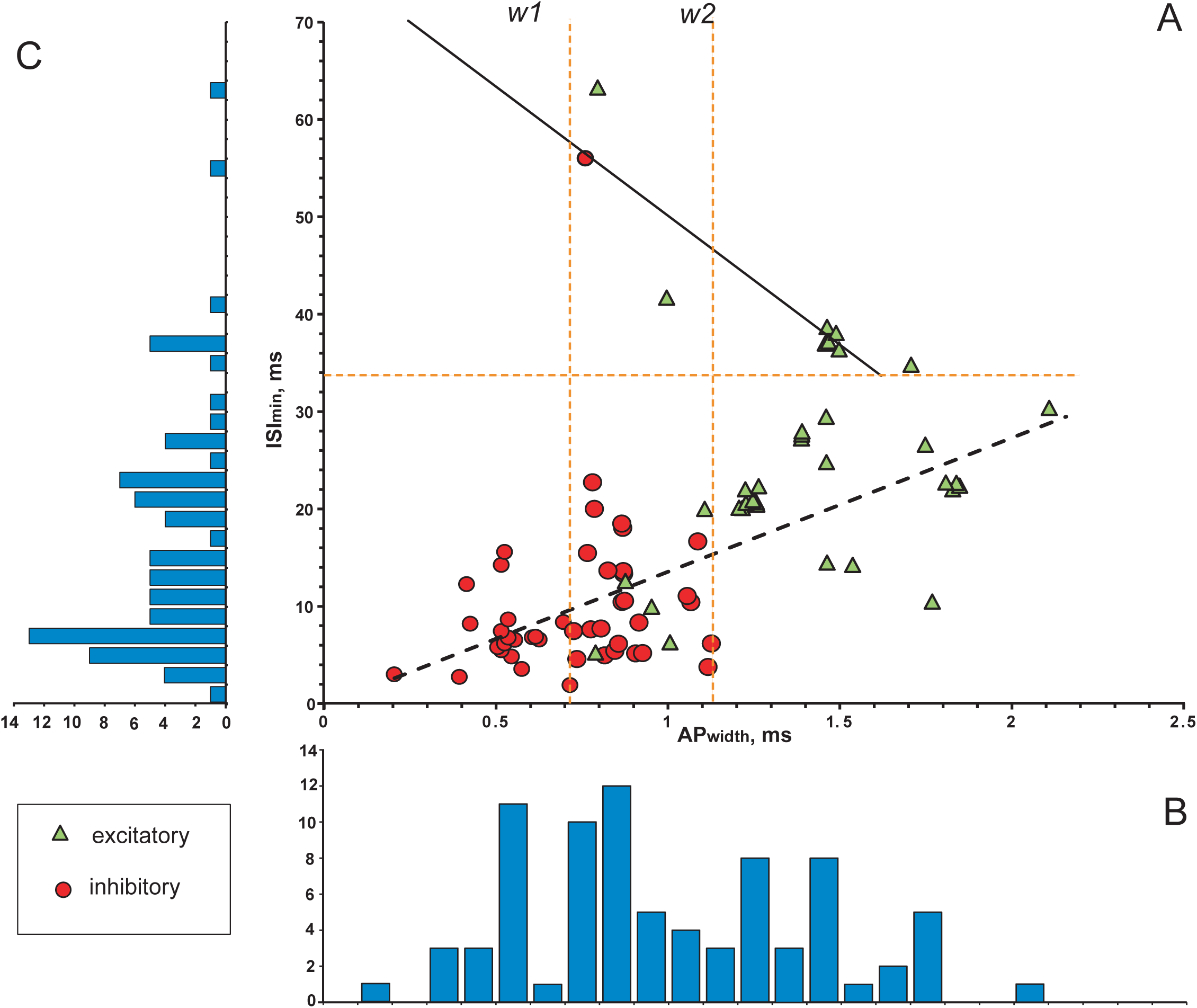
Relationships between the width of action potentials (AP_width_) and minimum of inter-spike intervals (ISI_min_) for 84 neuron types and subtypes. **A.** AP_width_ - ISI_min_ scatter diagram with results of linear regression. Green triangles and red circles indicate excitatory and inhibitory neurons, respectively. Dashed orange lines: horizontal line separates neurons with slow spikes from neurons with fast and moderate spikes; vertical lines (*w1* and *w2*) separate neurons with narrow, medium and wide action potentials. Black lines: solid line shows linear fitting for slow spike neurons with a function Y = - 26.72X + 76.42 (R^2^=0.83); dashed lie shows general linear fitting for fast and moderate spike neurons with a function Y = 13.79X - 0.05 (R^2^=0.52). **B**. AP_width_ histogram. **C**. ISI histogram.

Among the 22 neuron types/subtypes from the group with AP_width_<0.72 ms, 13 demonstrated so-called fast spiking behavior, which is distinguished by narrow spikes, high firing rate, and the absence or weak expression of spike frequency adaptation (Jonas et al., 2004). Besides these common characteristics, however, their firing patterns vary broadly even from a qualitative standpoint. Five of these 13 neuron types belong to the PSTUT family, namely CA3 Trilaminar (Gloveli et al., 2005), CA3 Aspiny Lucidum ORAX (Spruston et al., 1997), CA2 Basket (Mercer et al., 2007), CA1 Axo-axonic (Pawelzik et al., 2002), and CA1 Radial Trilaminar (Tricoire et al., 2011). Three types belong to the NASP family: DG Basket (Savanthrapadian et al., 2014), CA1 Horizontal Axo-axonic (Tricoire et al, 2011), and MEC LIII Superficial Multipolar Interneuron (Kumar and Buckmaster 2006). Two types, CA3 Axo-axonic (Dugladze et al., 2012) and CA2 Bistratified (Mercer et al., 2007), belong to the simple adapting spiking family; two types, DG HICAP (Mott et al., 1997) and DG AIPRIM (Lubke et al, 1998; Scharfman 1992), belong to the ASP-NASP family; and lastly CA1 Basket (Lee et al., 2011) belongs to non-persistent stuttering family.

Additionally, firing pattern families are unequally distributed among the groupings revealed by the above analysis. Persistent and non-persistent stuttering families and non-persistent bursting phenotypes are composed entirely of neuron types with narrow and medium fast/moderate spikes. Conversely, the rapidly adapting – non-adapting spiking phenotype is represented solely by neurons with spikes of intermediate width.

## Discussion

Neurons differ from each other by morphological and molecular features including the diversity and distribution of ion membrane channels in somata and dendrites. These intrinsic properties determine important physiological functions such as excitability, efficacy of synaptic inputs (Häusser et al., 2000; London et al., 2002; Komendantov and Ascoli, 2009), shapes of individual action potentials and their frequency (Bean, 2007), and temporal patterns (Mainen and Sejnowski, 1996; Krichmar et al., 2006).

In the neuroscience literature, the firing patterns of neuronal activity are commonly used to characterize or identify groups of neurons. Examples include descriptions of “strongly adapting, normally adapting, and nonadapting cells” (Mott et al., 1997); “fast-spiking and non-fast-spiking” interneurons (Bjorefeldt et al., 2016); “late spiking” cells (Tamas et al., 2003); “stuttering interneurons” (Song et al., 2013); “bursting” and “non-bursting” neurons (Hablitz and Johnston, 1981; Maskawa et al., 1982); “regular spiking, bursting, and fast spiking” (McCormick et al., 1985), and many more. However, it has until now remained challenging to integrate these characterizations across different laboratories and studies besides largely qualitative summaries.

In this study, we show that a quantitative, data-driven methodology based on the analysis of transients and steady states of evoked spiking activity can meaningfully classify the firing patterns of hippocampal neuronal types. This work is a further development of the effort initiated by the Petilla Interneuron Nomenclature Group (2008), which was applied to firing patterns in cortical neurons (Druckmann et al., 2013; Markram et al., 2015). At the same time, this work demonstrates the feasibility of systematic, comprehensive meta-analysis of electrophysiological data from the published literature. This is especially important as a necessary approach to help link and interpret the growing information from centralized, large-scale, “industrial” neuroscience projects (Kandel et al., 2013; Migliore et al., 2018; Teeter et al., 2018), with the distributed accumulation of data in traditional research laboratories (Ferguson et al., 2014).

From the electrophysiological recordings of 90 neuron types in the rodent hippocampus, we identified 23 firing patterns, 15 of which were completed, that is, included both transient(s) and putative steady state components (see Figs. 4 and 5). Taking into consideration the firing pattern information enables a possible refinement of neuron type delineation by identifying 52 putative electrophysiological subtypes among 22 neuron types. Subsequent two-step cluster analysis allows for the clear distinguishing of 9 unique families of 44 firing pattern phenotypes among 120 neuron types and putative subtypes. Notwithstanding the focus of the present research on the hippocampal formation, the firing pattern classification framework introduced with this study can be readily applied to spiking activity of neurons from other brain regions.

The two firing pattern families characterized by bursting phenotypes (transient and persistent) are comprised of excitatory neurons, while the persistent stuttering family only included inhibitory neurons. However, the majority of phenotype families are mixed between putatively glutamatergic and GABAergic types (Fig. 6B). Thus, the identification of a firing pattern phenotype by itself is a useful but in most cases insufficient attribute for a reliable categorization of excitatory and inhibitory neurons.

The frequency of discharges is an important characteristic of neuronal communication. Many neuron types, especially interneurons, show fast spiking behavior: they are capable of firing at high frequencies (200 Hz or more) with little decrease in frequency during prolonged stimulation (Jonas et al., 2004; Bean 2007). Spike frequency correlates with electrophysiological characteristics, such as action potential duration or fast AHP amplitude (Druckmann et al., 2013). Fast spiking neurons typically have narrow action potentials and high-amplitude fast AHP (Bean 2007). Our correlation analysis of Hippocampome.org data reveals that transient stuttering (TSTUT.) is not typical for cells with extremely high-amplitude fast AHPs and delayed firing (D.) is not characteristic for neuron types with wide action potentials (Box 1). Interestingly, plotting ISI_min_ against AP_width_ for all neuron types with relatively faster firing (maximum frequencies higher than ∼30 Hz) and for all neuron types with slower firing (maximum frequencies lower than 29 Hz) reveals opposite, statistically significant linear relationships (Fig. 8A).

Firing pattern phenotypes of central mammalian neurons are determined by biophysical properties associated with expression and distribution of several types of Ca^2+^ and K^+^ channels, which modulate specific ion currents (Llinás 1988; Migliore and Shepherd, 2005; Bean, 2007), as well as with expression of other molecular markers (Caballero et al., 2014; Markram et al., 2004; Petilla Interneuron Nomenclature Group et al., 2008). Despite the relative sparsity of molecular marker information, analysis of the correlations between firing patterns and other neuronal properties revealed novel interesting relationships in hippocampal neuron types (see Box 1 for illustrative examples).

Firing patterns play important roles in neural networks including the representation of input features, transmission of information, and synchronization of activity across separate anatomical regions or distinct cell assemblies. Although single spikes can provide temporally precise neurotransmitter release, this release usually has low probability in central synapses. Neurons can compensate for the unreliability of their synapses by transmitting signals via multiple synaptic endings or repeatedly activating a single synapse (Lisman, 1997). Thus, spikes grouped together in bursting or stuttering activity increase the probability of transmission via unreliable synapses compared to separate spikes with the same average frequency. In the hippocampus, a single burst can produce long-term synaptic potentiation or depression (Lisman, 1997). It has also been hypothesized that, due to the interplay between short-term synaptic depression and facilitation, bursting with certain values of ISIs are more likely to cause a postsynaptic cell to fire than bursts with higher or lower frequencies (Izhikevich et al., 2003). Recent results have also revealed that single bursts in hippocampal neurons may selectively alter specific functional components of the downstream circuit, such as feedforward inhibitory interneurons (Neubrandt et al., 2018).

Experimental studies provide strong evidence that different brain circuits employ distinct schemes to encode and propagate information (Xu et al, 2012): while information relay by isolated spikes is insignificant for the acquisition of recent contextual memories in the hippocampus, it is essential for memory function in the medial prefrontal cortex. However, even within the hippocampus, different neuronal circuits may employ distinct coding schemes by relying on isolated spikes or bursts of spikes for execution of critical functions (Xu et al, 2012). Indeed, distinct sub-regions of the hippocampal formation show differential distributions of spiking, bursting, and stuttering firing pattern phenotypes (Fig. 6).

In this study, the phenotyping of most types of neurons was relied on the digitization and quantitative analysis of single (or limited numbers of) experimental recordings of electrical activity extracted from many relevant publications. Until neuroscience switches to the systematic deposition of all firing traces recorded and analyzed for a given publication to public repositories, such representative illustrations, however limited, constitute a fairly accurate reflection of the communal knowledge about neuronal physiology in particular neural system. Thus, our approach is based on the statistical quantification of integrated data presented in the literature.

The findings presented in this report resulted from the analysis of firing patterns in response to depolarizing current. To this date, this is by far the most common experimental protocol for characterizing the neuronal input-output function. Nevertheless, different types of neurons also exhibit distinct responses to hyperpolarization, as well as to its termination. For example, several neuron types described in Hippocampome.org demonstrate rebound spiking: CA1 Trilaminar (Tricoire et al., 2011, Sik et al., 1995), CA1 Back-Projection (Sik et al., 1994), CA1 O-LM (Sik et al., 1995), CA1 SO-SO (Pawelzik et al., 2002), MEC LIII Multipolar Interneuron (Kumar and Buckmaster 2006), MEC LII Stellate (Canto and Witter 2012b), MEC LII Oblique Pyramidal (Canto and Witter 2012b). Such neuronal behaviors, owing to the hyperpolarization-activated cation current (h-current), may play an important role in hippocampal rhythmogenesis (raHasselmo 2014) and could be locally modulated by activity-dependent changes in intrinsic excitability (Ascoli et al., 2010). It will therefore be interesting to extend the current firing pattern phenotyping by considering these additional neuronal properties in future work.

The information on firing patterns of neuron types further expands the rich knowledge base of neuronal properties Hippocampome.org, which already contained information on morphology, molecular marker expression, connectivity, and other electrophysiological characteristics (Wheeler et al., 2015). Computation of the potential connectivity map of all known 122 neuron types by supplementing available synaptic data with spatial distributions of axons and dendrites enabled the reconstruction of a circuitry containing more than 3200 putative connections (Rees et al., 2016).

Further development also includes simulation of firing activity of different neuron types based on dynamical systems modeling (Venkadesh et al., 2018). This ongoing accumulation of data and knowledge makes Hippocampome.org a powerful tool for building real-scale models of the entire hippocampal formation, thus substantially expanding the potential scope of recent advances in this regard (Bezaire et al., 2016). More generally, such knowledge bases are playing an increasingly important role in neuroscience research by fostering computational analyses and data-driven simulations.

---

**Table 1. Abbreviations:** Pct. – percentage of firing pattern recordings for which this solution was used.

**Table 2. Abbreviations:** KAc – potassium acetate (KCH_3_COO); KGlu – potassium gluconate; KMeSO_4_ – potassium methylsulphate (CH_3_KSO_4_); PCr – phosphocreatine; Pct. – percentage of firing pattern recordings for which this solution was used; 10 mM HEPES (4-(2-hydroxyethyl)-1-piperazineethanesulfonic acid) was used in all patch pipette solutions. Asterisks indicate examples of micropipette solutions.

**Table 3. Abbreviations:** *a*_*1*_ – slope of linear fitting for normalized ISIs *vs* normalized time; *DF* – delay factor; *f*_min_ – minimum frequency of stuttering or bursting; *F*_*pre*_, *F*_*post*_, *F*_*PSTUT*_, *F*_*PSWB*_ – ISI comparison factors, *ISI*_*max*_ – maximum inter-spike interval; *p*_2,1_ – *p*-value for differences between two-and one-parameter linear fitting; *p*_3,2_ – *p*-value for differences between three-and two-parameter linear fitting; *PFS* – post firing silence; *SF* – silence factor; *S*_*RASP*_ – slope of linear fitting of rapid transient; *SWA* – slow wave amplitude; *SWA*_min_ – minimum slow wave amplitude.

**Table 4. NASP** – HICAP (Mott et al. 1997, Fig. 11A); **PSTUT** - CA1 Neurogliaform (Fuentealba et al. 2010, Fig.5B); **PSWB** - CA3 Pyramidal (Bilkey and Schwartzkroin 1990, Fig. 1a); **ASP.NASP** - CA3 Basket-CCK (Gulyás et al. 2010, Fig. 1b, right); **ASP.SLN** – EC MEC LV Pyramidal (Canto and Witter 2012b, Fig.10C7); **RASP.NASP** – EC LV Deep Pyramidal (Hamam et al. 2000, Fig.3C); **RASP.SLN** – CA1 Radiatum Giant (Bullis et al. 2007, Fig.5A); **TSTUT.NASP** - EC LV Deep Pyramidal (Hamam et al. 2002, Fig.5E); **TSTUT.PSTUT** - CA1 (Price et al. 2005, Fig.3A2); **TSUT.SLN** – CA2 SP-SR (Mercer et al. 2012; Fig. 3A); **TSWB.NASP** - CA1 Pyramidal (Zemankovics et al. 2009, Fig. 1B);**TSWB.SLN** - CA3 Pyramidal (Hemond et al. 2008, Fig. 4); **D.NASP** – DG Neurogliaform (Armstrong et al. 2011, Fig.3A, top trace); **D.PSTUT** - CA2 Basket (Mercer et al. 2007, Fig. 5B)**; D.PSWB -** cultured *rutabaga* mutant giant neuron of *Drosophila* (Zhao and Wu 1997, Fig.7, *top left*); **D.ASP.SLN -** neuron in external lateral subnucleus of lateral parabrachial nucleus (Hayward and Felder 1999, Fig.3A, top); **D.RASP.NASP** - CA3 LMR-Targeting (Ascoli et al. 2009, Fig. 1A); **D.TSUT.SLN** - striatal fast-spiking neuron (Sciamanna and Wilson 2011, Fig. 1C); **D.TSWB.NASP** - CA1 Axo-Axonic (Buhl et al. 1994, Fig. 5D). **Abbreviations:** Lat. – lateral; nucl. – nucleus.

## Acknowledgements

This work was supported by grants from the National Institutes of Health (NS39600) and the National Science Foundation (IIS-1302256). We thank Charise M. White and Keivan Moradi for useful discussions and Amar Gawade for help with the web portal. The authors declare no competing financial interests.

